# Life out of water: Genomic and physiological mechanisms underlying skin phenotypic plasticity

**DOI:** 10.1101/772319

**Authors:** Yun-wei Dong, Tessa S. Blanchard, Angela Noll, Picasso Vasquez, Juergen Schmitz, Scott P. Kelly, Patricia A. Wright, Andrew Whitehead

## Abstract

The Devonian radiation of vertebrates from aquatic into terrestrial habitats required behavioral, physiological, and morphological adaptations. Changes to skin structure and function were likely crucial, but adaptations were needed to resolve contrasting demands of maintaining a mechanical and physiological barrier while also facilitating ion and gas transport. Little is known of the mechanisms that underlie skin plasticity and adaptation between water and air. We performed experiments using two isogenic lineages of an amphibious killifish (*Kryptolebias marmoratus* from brackish and freshwater habitats) and used transcriptional and morphological data to reveal mechanisms recruited to resolve the dual challenges of skin providing both a barrier and an exchange interface during terrestrial acclimation. Transcriptional regulators of skin morphogenesis were quickly activated upon emersion. Regulation of cell-cell adhesion complexes, coupled with pathways homologous with those that regulate stratum corneum formation, was consistent with barrier function and mechanical reinforcement. Cutaneous respiration was associated with regulation of angiogenesis pathways and with blood vessel architecture that facilitated extremely short diffusion distances and direct delivery to ionocyotes. Evolutionary analyses revealed directional selection operating on proteins involved in barrier and respiratory functions, reinforcing the importance of these mechanisms for enabling the amphibious lifestyle of *K. marmoratus*. Fish from brackish niches were more resilient to emersion and also differed from freshwater fish in ionoregulatory responses to emersion. We conclude that plasticity of barrier, respiratory, and ionoregulatory functions in skin evolved to support the amphibious lifestyle of *K. marmoratus*; similar processes may have facilitated the terrestrial radiation of ancient fishes.

**Significance statement:** The transition of vertebrate life from water to land coincided with solving multiple physiological challenges including avoiding drying out while also exchanging gases and ions with the environment. Though changes in the skin were likely important, little is known of the mechanisms that underlie skin flexibility and adaptation between water and air. We performed air exposure experiments with an amphibious killifish; gene expression profiling, microscopy, and evolutionary analysis of proteins revealed cell structures, proteins, and molecular pathways that support skin flexibility and adaptations during air exposure, and ion regulation contributed to differences in killifish abilities to adjust to air. Amphibious killifish are useful models to help us understand changes that enable water to air transitions in contemporary and ancient fishes.

## Introduction

Devonian fishes seeded the first successful vertebrate invasion of land, where various phenotypic innovations promoted a dramatic terrestrial radiation. What were the mechanistic changes that facilitated this crucial adaptive leap to life on land? It is difficult to infer the genetic and physiological traits that enabled the original Devonian invasion of land. However, modern fishes have repeatedly evolved physiological and behavioral traits that enable exploitation of terrestrial niches (1). Extant amphibious fishes offer an opportunity to study the variety of mechanisms that may have facilitated the challenging transition of vertebrate lifestyles between aquatic and terrestrial habitats.

Transition between aquatic and terrestrial habitats involves multiple phenotypic traits. Perhaps best studied are changes in skeletal morphology, at least partly because these traits can be preserved in the fossil record (2). However, other morphological and physiological changes, which are less well preserved in fossils, are also necessary to enable terrestrial adaptation. For example, it would be crucial to change the structure and function of: respiratory organs in order to support aerial respiration, water-exposed barrier tissues to prevent desiccation during emersion, and ion regulatory/excretory organs to assist in the maintenance of osmotic homeostasis as well as facilitate nitrogenous waste excretion in air (3). In this regard, the opposing pressures of life in both water and land have been studied in amphibians (4), where developmental changes during metamorphosis allow for water to land transition. But the mechanisms that resolve the opposing pressures of an amphibious existence in fishes have received comparatively less attention. Nevertheless, this is an area worthy of investigation because the skin of amphibious fishes is broadly acknowledged to modulate in response to all of the aforementioned challenges associated with the water to air transition, and life out of water for an amphibious fish is more likely to resemble the challenges faced by terrestrial tetrapods during a pivotal period in vertebrate evolution.

For the skin of amphibious fishes to become a crucial organ for oxygen transport, ion and water regulation, and nitrogenous waste excretion, and to provide a mechanical barrier out of water, we hypothesized that structural, physiological, and molecular changes must occur soon after emersion. We therefore predicted that rapid and sustained regulation of molecular pathways associated with dermal/epidermal structural remodeling, gas exchange, ammonia detoxification, and water and ion transport, underpin the alterations in skin structure and function that unfold during this transition from immersion to emersion.

To test these hypotheses we used the amphibious mangrove rivulus (*Kryptolebias marmoratus*), which is a particularly opportune model system for studying the physiological and genomic mechanisms that support terrestrial acclimation. The species is native to the Western Atlantic mangrove forests, from southern Florida to Brazil. They frequently escape water for terrestrial microhabitats, including the mud walls of crab burrows, damp leaf litter, or rotting logs, for variable durations up to 66 days (5).

We compared two clonal strains of *K. marmoratus* during acclimation from water to air. One lineage is native to brackish water habitats in Honduras (BW) and the other to freshwater habitats in Belize (FW). Lineages were independently perpetuated by self-fertilization resulting in a purging of within-lineage genetic variation and creation of isogenic strains (6). Strains were acclimated to their native salinities, and then over the course of one week following emersion we measured changes in survival, metabolic rate, skin structure (light microscopy, transmission electron microscopy), and skin genome-wide gene expression (RNA-seq) to infer skin mechanisms that resolve the multiple competing physiological demands of terrestrial acclimation. We predicted that genotype and osmotic acclimation would interact to provoke between-strain differences in the regulation of some of these mechanisms, and thereby provide additional insight into mechanisms that underlie variable terrestrial acclimation abilities. Finally, we used models of comparative protein evolution to identify genes evolving by directional selection in *K. marmoratus* compared to other fish species that have limited or no terrestrial acclimation ability. We predicted that genes evolving by directional selection uniquely in the *K. marmoratus* lineage will be associated with the molecular pathways that regulate the species’ unusual terrestrial acclimation abilities.

## Methods

### Fish collection and maintenance

Two hermaphroditic strains of *Kryptolebias marmoratus* were maintained individually in the Hagen Aqualab at the University of Guelph, Ontario. A strain originally collected from Honduras (HON) in 1996 was held in the lab (15 ppt, 25°C) for 20 years (∼60 generations; HON11; (6)). A freshwater strain (FW) captured in Dec. 2012 at Long Caye, Belize in an open pool (0.3 ppt) was held in the lab (0.3 ppt, 25°C) for 3.5 years (∼10 generations; (7)). The propagation of self-fertilizing hermaphroditic *K. marmoratus* for at least 10 generations results in inbred stocks of isogenic individuals (6). Fish were held singly in plastic containers (120 ml Fisherbrand; Fisher Scientific) in 60 ml of water under constant conditions (12:12 h light:dark cycle, 25°C; (8)). Reverse osmosis water and sea salt (Instant Ocean TM, Crystal Sea, Baltimore, MA, USA) were used to formulate water to the appropriate salinity and changed weekly. Fish were fed live *Artemia* three times a week. The University of Guelph Animal Care Committee approved this project (AUP 2239).

### Emersion challenge experiment

HON (0.120g ± 0.005g) and FW (0.130g ± 0.006g) fish were acclimated to air for up to 7 days at 25°C. Before aerial transfer, tissues from immersed fish were sampled as a pre-transfer (pre-emersion) control (0 hours). Upon aerial transfer (emersion challenge) fish were maintained on moist filter paper (15 ppt or 0.3 ppt) in plastic rearing containers as previously described (9) and skin samples were collected at 1 hour, 6 hours, 24 hours (1 day), 72 hours (3 days), and 168 hours (7 days) for transcriptomics. Fish were euthanized with tricaine methanesulfonate (MS-222; 1.5mg/mL). Skin was dissected from the mid-region of the body by making a ventral incision and carefully removing extraneous tissue and muscle from the skin, then stored in RNAlater (Thermo Fisher Scientific, Markham, ON, Canada) and archived at −20°C. For histology, fish were acclimated to water or air (0h and 168h), euthanized by immersion in ice-cold water, then fixed and preserved for either collagen staining or transmission electron microscopy (TEM) imaging.

### Physiological profiling

Survival was measured using a different set of fish (HON, n=20; FW, n=20) that were air-exposed for up to 7 days and monitored daily. The experiment was terminated if survival reached 20% as in the case of the FW fish. Routine metabolic rate (µmol O2·g-1·h-1) was measured as described previously (8). Briefly, fish were placed in glass respirometry chambers (∼1 ml), allowed to recover from handling stress and changes in air oxygen levels were measured over a 10-minute period under closed conditions at 25°C (Loligo Systems WITROX 4, Tjele, Denmark). Optodes were calibrated weekly using air (100% DO) and sodium sulfite (2M; 0% DO). Treatment effects on metabolic rate were tested using two-way ANOVA, with time and strain specified as main effects, including an interaction term, followed by a *post hoc* Holm–Šidák test.

### Profiling of skin structure

For collagen staining, fish were fixed overnight (4°C) in 10% buffered formalin, decalcified (Cal-Ex, Fisher Scientific) for one hour (room temperature), and stored in 70% ethanol (4°C) until embedding in paraffin wax. Tissues were sliced in a transverse orientation (5µm), deparaffinized in xylene, and rehydrated in an ethanol graded series (10). Tissues were then stained with picrosirius red, a collagen-specific dye (Electron Microscopy Sciences, Hatfield, PA, USA), and photographed under circularly polarized light as previously described (11). The mean collagen hue for each picture was calculated according to the pixel distribution as previously described (12). Treatment effects on collagen staining were tested using two-way ANOVA, with time and strain specified as main effects, including an interaction term, followed by a *post hoc* Holm–Šidák test.

Samples for TEM observation were prepared using skin sampled from both HON and FW animals that were either pre-emersion controls or had been exposed to emersion challenge for 7 days. Samples were acquired from a standardized region of skin, just posterior to the operculum and dorsal of the horizontal midline. Tissue was removed to include a small section of underlying epaxial muscle so as not to damage the skin, and to provide orientation when sectioning. Isolated tissue was placed immediately into freshly prepared, ice-cold glutaraldehyde fixative (2.5% glutaraldehyde in 0.1M sodium cacodylate buffer, pH 7.3). Samples were fixed at 4 °C overnight and then rinsed twice in 0.1 M sodium cacodylate buffer (2 x 10 min, pH 7.3). Samples were post-fixed in osmium teroxide (1% osmium teroxide in 0.1M sodium cacodylate buffer, pH 7.3), rinsed twice in buffer, and dehydrated through a graded ethanol series (70%, 90%, and 3 x 100%, 10 min each), then immersed in propylene oxide (2 x 10 min), and transitioned into Quetol-Spurr resin (2 h in 1:1 propylene oxide/Quetol-Spurr resin followed by 2 h in Quetol-Spurr resin). Final infiltration in Quetol-Spurr resin was carried out overnight, after which sample pieces were placed in embedding molds and polymerized at 65 °C overnight.

Sections (90nm) were prepared using a Leica EM UC7 ultramicrotome, stained with uranyl acetate and lead citrate, and viewed in an FEI Tecnai 20 TEM. Skin morphometric analysis was conducted using Image J software (https://imagej.nih.gov/ij/). Sub-apical desmosome #2 was the second desmosome observed in the sub-apical region of the epidermis and its depth was determined by tracing its length. On average, for each fish, the lengths of typically 6 – 10 desmosomes were measured and then averaged to give an N of 1 for that animal. A total of 158 desmosomes were measured for 20 animals. Treatment effects on desmosome length were tested using two-way ANOVA, with time and strain specified as main effects, including an interaction term.

### Detecting positive selection

Positively selected molecular changes that are lineage-specific underlie adaptive phenotypes that may be unique to a species. They leave recognizable footprints in the genome, for example in the amino acid sequence of proteins. Such footprints are detected when sequences are compared between orthologous loci among relatives. We used OrthoMCL (13) which is a program that filters orthologous proteins and GUIDANCE2 (14) to remove unreliable coding sequence (CDS) alignments and alignment positions. We then used Phylogenetic Analysis by Maximum Likelihood (PAML (15)) to identify positively selected sites on targeted lineages, in our case those that are specific for *K. marmoratus* (Fig. S15).

We downloaded protein and CDS sequences in FASTA format of genome assemblies from (1) *K. marmoratus* (mangrove rivulus), (2) *Austrofundulus limnaeus* (bony fishes), (3) *Nothobranchius furzeri* (turquoise killifish), (4) *Poecillia formosa* (Amazon molly), (5) *Fundulus heteroclitus* (mummichog), (6) *Oryzias latipes* (Japanese medaka; ASM223467v1), (7) *Gasterosteus aculeatus* (three-spined stickleback), (8) *Takifugu rubripes* (fugu), and (9) *Danio rerio* (zebrafish) from NCBI and the Broad Institute (Fig. S15). To assign a database of orthologous sequences from these nine sets of genome protein FASTA sequences, we used OrthoMCL v2.0.9 (13), and the following criteria were applied to achieve optimal clustering resolutions: similarity *p*-value ≤ 1e-05, protein percent identity ≥ 40%, and MCL inflation of 1.5. In a preliminary clustering step, compliant FASTA files were established by applying the module orthomclAdjustFasta. The OrthoMCL module orthomclFilterFasta was used to remove proteins of poor quality to finally generate the goodProtein.fasta-file compiling all proteins (337,958 proteins; 230,1 MB) with specific assigned headers for the subsequent all-vs-all BLAST.

To build orthologous groups, we first conducted an all-vs-all NCBI BLAST (ncbi-blast-2.2.24+). With the OrthoMCL module orthomclBlastParser, similarities were then summarized within a file that was loaded to mysql by the orthomclLoadBlast module. The orthomclPairs and orthomclDumpPairsFiles modules then found, sorted, and created results files of all protein pairs (orthologs, in-paralogs, co-orthologs). The mcl and orthomclMclToGroups modules generated the groups.txt-file. We then used a custom python script to extract only unique genes that were represented a) in all species, and b) due to their critical phylogenetic distance and poorly sequenced genomes, in all species except *D. rerio* and/or *T. rubripes* and/or *G. aculeatus*. We then extracted all corresponding nucleotide sequences from the CDS FASTA file by applying another custom python script. GUIDANCE2 was then used to remove unreliably aligned positions in the multiple sequence alignments before testing for positively selected sites with PAML 4 (15). We conducted the analysis twice (e.g., the GUIDANCE2 filter and subsequent PAML analysis) using different aligners: a) Muscle version 3.8.31 (16) and b) Mafft version X7.721 (17). Figure S15 lists all guide trees that were used to fulfill branch-site tests for positive selection (M2-BS-H1 vs M2-BS-H0) in the *K. marmoratus* lineage using PAML. We inferred positive selection in *K. marmoratus* when the alpha-value for the likelihood ratio test was <0.01.

### Transcriptome change during emersion acclimation

Transcriptomics data collection and analyses were described in (8). Briefly, tissues were homogenized and RNA extracted using RNAeasy purification kits (Agilent, Inc.). Individually-indexed RNA-seq libraries were prepared using NEBnext RNA library preparation kits for Illumina, libraries were pooled, and sequence data collected across four lanes of Illumina 4000 (PE-150). Five biological replicates were included per treatment group, except for two treatments (HON11 fish 72hrs post-emersion, and HON11 fish 0hrs pre-emersion control) that had four replicates. Sequence reads were mapped to the *K. marmoratus* gene set reported in (18) (Whole Genome Shotgun project GenBank accession LWHD00000000, GenBank assembly accession: GCA_001649575.1) using STAR v2.4.2a (19), and read counts generated using HTseq v0.6.1 (20). We retained genes that had greater than 10 read counts for at least four of five biological replicates within any treatment group. Read counts were log2 transformed and normalized for gene length and total library size in EdgeR v3.20.9 (21). Differential expression analysis was performed in Limma v3.34.9 (22), where *strain* (HON, FW)) and *time* (0hr, 1hr, 6hrs, 24hrs, 72hrs, 168hrs) were specified as main effects, and *strain*\**time* as an interaction term, including false discovery rate adjustment of p-values. RNA-seq read sequences are archived at NCBI (SRA accession: SRP136920, BioProject: PRJNA448276). A matrix of per-gene per-sample raw read counts (“final.out.txt”) and R scripts that detail all of these analyses are available in a GitHub repository (https://github.com/WhiteheadLab/mangrove_killifish).

We examined three sets of transcriptional responses: transcripts with a significant temporal effect that was similar between HON and FW fish (shared acclimation response genes), transcripts with a significant time-by-strain interaction (genes that differed in their acclimation response between HON and FW fish), and transcripts that were not responsive to emersion, but were constitutively differentially expressed between HON and FW fish (strain genes). Shared acclimation response genes were grouped by temporal patterns of co-expression, which were defined by the time post-emersion at which expression was first significantly different from pre-emersion controls. This grouping was performed separately for up and down-regulated (compared to pre-emersion control) genes. For functional inference, clusters of co-expressed genes were examined for gene ontology (GO) and KEGG pathway enrichment using DAVID v6.8 (23) (Tables S1-S11) using Uniprot IDs from human orthologs of *K. marmoratus* genes, and for canonical pathway and BioFunction enrichment using Ingenuity Pathway Analysis (24).

## Results & Discussion

Though the HON and FW animals shared many molecular, morphological, and physiological responses during aerial acclimation, they differed markedly in their resilience to emersion. All HON animals survived the entire duration of emersion, whereas FW animals suffered 80% mortality by 7 days out of water (Fig. 1A). Variable resilience between strains could be due to genetic differences between strains, or due to differences in acclimation environment. Though the FW strain used in these experiments inhabit a very different native habitat (freshwater) than HON strain animals (brackish/marine), likely resulting in evolved differences in physiology, the experiments reported here were not designed to distinguish genetic (heritable) from acclimation differences between strains. By 7 days following emersion metabolic rate decreased by 5.2% in HON animals, but not in FW animals (time-by-strain interaction p = 0.02; Fig. 1B). Metabolic depression is likely an adaptive response to diminished feeding during terrestrial forays (Turko et al. in review) and is observed in at least two other brackish/marine strains of *K. marmoratus*, each of which also survived at least 7 days of emersion (8). High mortality following emersion is therefore a response that is confined to the FW strain of animals. We conclude that acclimation or adaptation to fresh water constrains effective acclimation to emersion.

**Figure 1.**
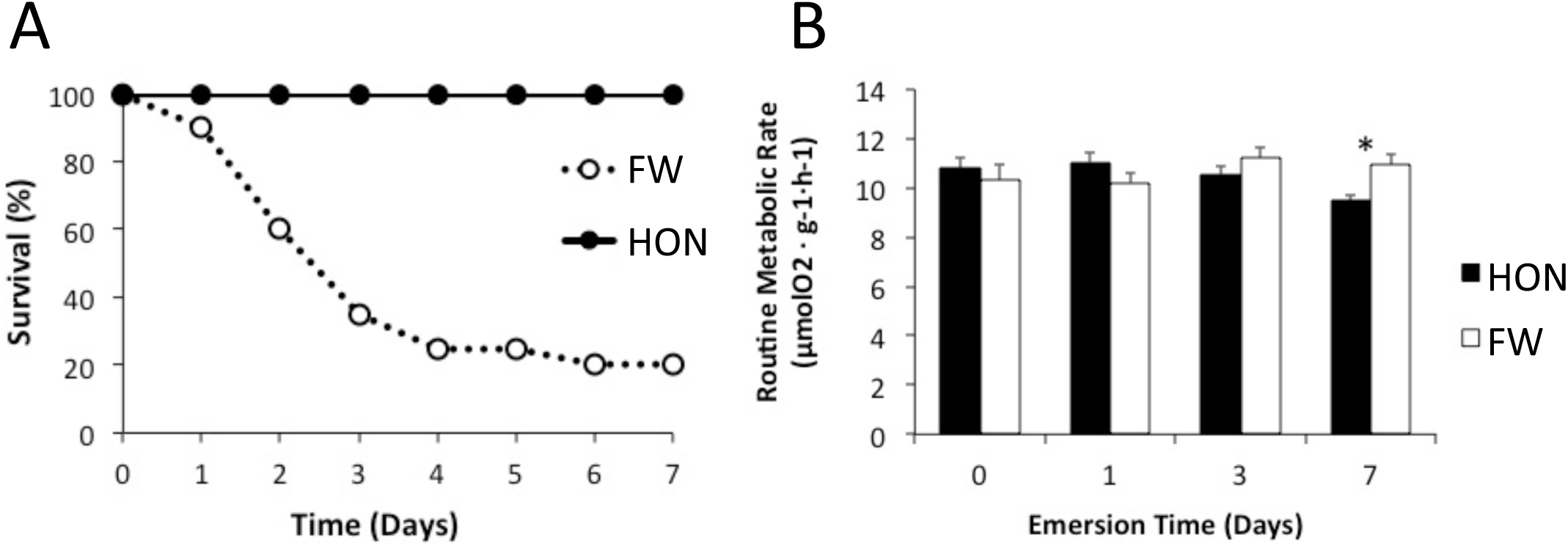
Survival (A) and routine metabolic rate (B; means ± s.e.m.) in water (time 0) and during a seven-day time-course of emersion for HON (closed black circles and bars) and FW (open white circles and bars) fish.

We detected many genes that were transcriptionally responsive to emersion, and that differed in expression between strains (Appendix S1). A large set of genes was transcriptionally responsive to emersion challenge in both strains (main effect of time only, with no significant time-by-strain interaction; Fig. 2A). When sorted by time point at which expression was first significantly different from pre-emersion controls, patterns of propagating waves of up- and down-regulation emerged. Up-regulation tended to be quick (early) and transient, whereas down-regulation took longer and tended to be more persistent. A smaller set of genes was differentially expressed between strains during acclimation (interaction between time and strain; Fig. 2B), and fewer still were constitutively differentially expressed between strains (main effect of strain only, Fig. 2C). Changes in expression of genes encoding structural proteins were further supported by changes in skin ultrastructure and collagen content in both strains.

**Figure 2.**
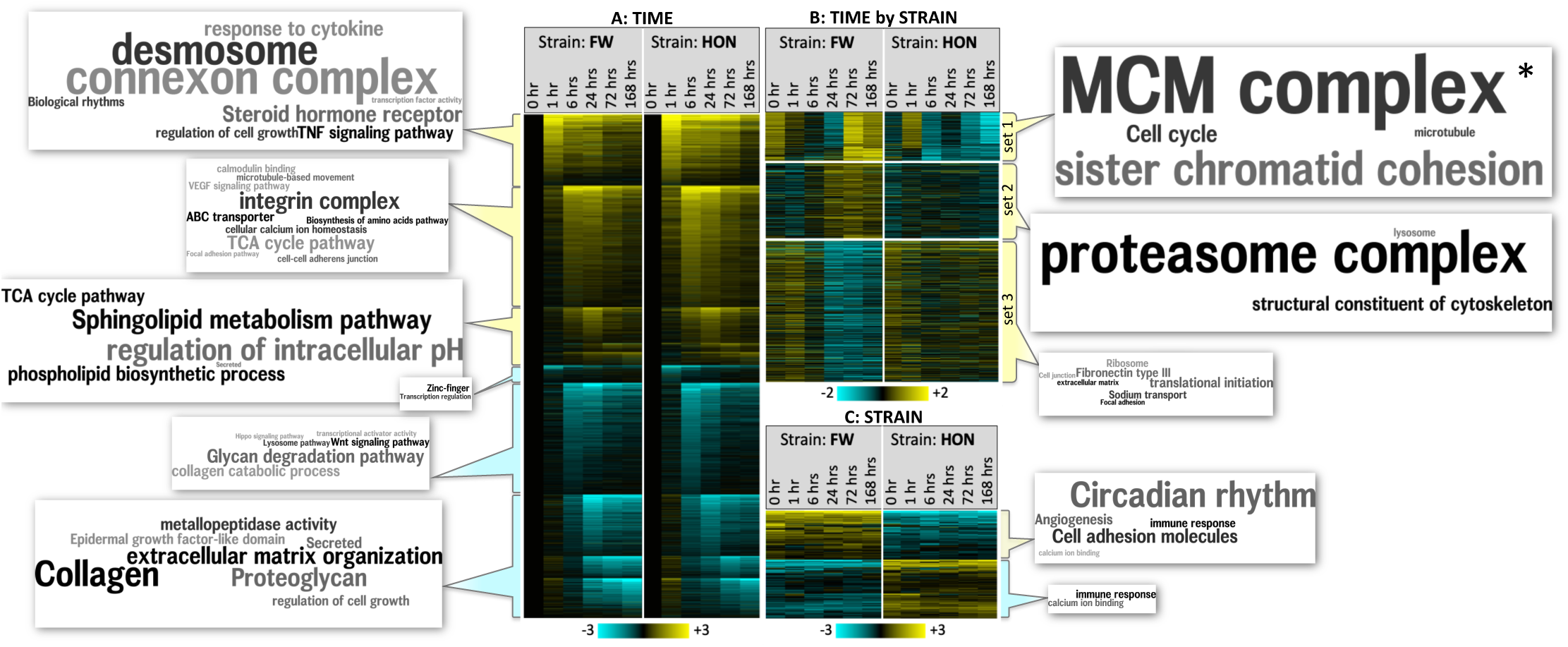
Functional enrichment of co-expressed sets of genes (word clouds), where heatmap color indicates log2 fold change in expression (yellow and blue indicate up- or down-regulation, respectively) during emersion relative to pre-emersion control (A) or relative to grand mean (B, C). A) Genes showing a main effect (p<0.001) of differential expression through time (hours), clustered by earliest time by which expression was significantly different from pre-emersion controls (0 hr) for both strains of fish (FW and HON). B) Genes showing a significant interaction between strain and time (p<0.05) clustered into three sets of co-expressed genes. C) Genes showing a main effect (p<0.05) of strain. Word clouds (www.wordle.net) represent significantly enriched (p<0.05) GO terms sampled from David clusters (Tables S1-S11), where font is linearly scaled by fold-enrichment of GO term (e.g., for genes upregulated by 1 hr post emersion, *transcription factor activity* is enriched 2.3-fold (p=4.5E-05), whereas *desmosome* is enriched 12.3-fold (p=6.1E-04); Tables S1-S11). Font size is normalized across all 11 word clouds. *fonts for this word cloud were down-scaled by 50% to fit within figure (e.g., *MCM complex* is enriched 36-fold).

### Skin morphology

Air exposure induced rapid changes in transcription of genes associated with skin remodeling. Transcription factors were enriched among the genes up-regulated by 1 hr post emersion (Table S1), including those known to regulate skin morphogenesis (25) such as transcription factors P65 (NFkB or RELA), retinoic acid receptor, Krueppel-like factor 4, and transcription factors AP-1 (JUN, FOS) (Fig. S1). Amphibian skin remodeling during metamorphosis is thyroid hormone-dependent; remodeling is initiated by thyroid hormone receptor which heterodimerizes with retinoic acid receptor and acts as a transcription factor to regulate the expression of target genes (4). We observed parallel up-regulation of THRA and RARG within 1 hr post-emersion (Fig. S1), which is consistent with a convergent role of thyroid hormone signaling in skin remodeling during terrestrial transitions between amphibians and amphibious fish (4, 26).

At cell junctions, multiple structures serve to attach epithelial cells together, in sheets, and to the basement membrane and extracellular matrix (ECM) (25), and regulate the exchange of small molecules and the passage of water. Desmosomes and adherens junctions are cadherin-containing complexes that form intercellular junctions providing strength to tissues, and focal adhesions are integrin-containing complexes that attach epidermal cells to the basement membrane and ECM. Furthermore, gap junctions (e.g., connexin-containing complexes) provide channels through which small molecules may pass between adjacent cells, and tight junction proteins (e.g., claudins, occludins) regulate paracellular permeability to water and small molecules. The molecular components of these structures, and the pathways that regulate them, are all transcriptionally regulated within hours of emersion (Figs. 2A, 3, S2–S5). Furthermore, TEM observations of *K. marmoratus* skin confirms that differences in the configuration of specialized epidermal structures associated with the functional integrity of the epidermis occurred either between HON and FW fish or following aerial acclimation.

**Figure 3.**
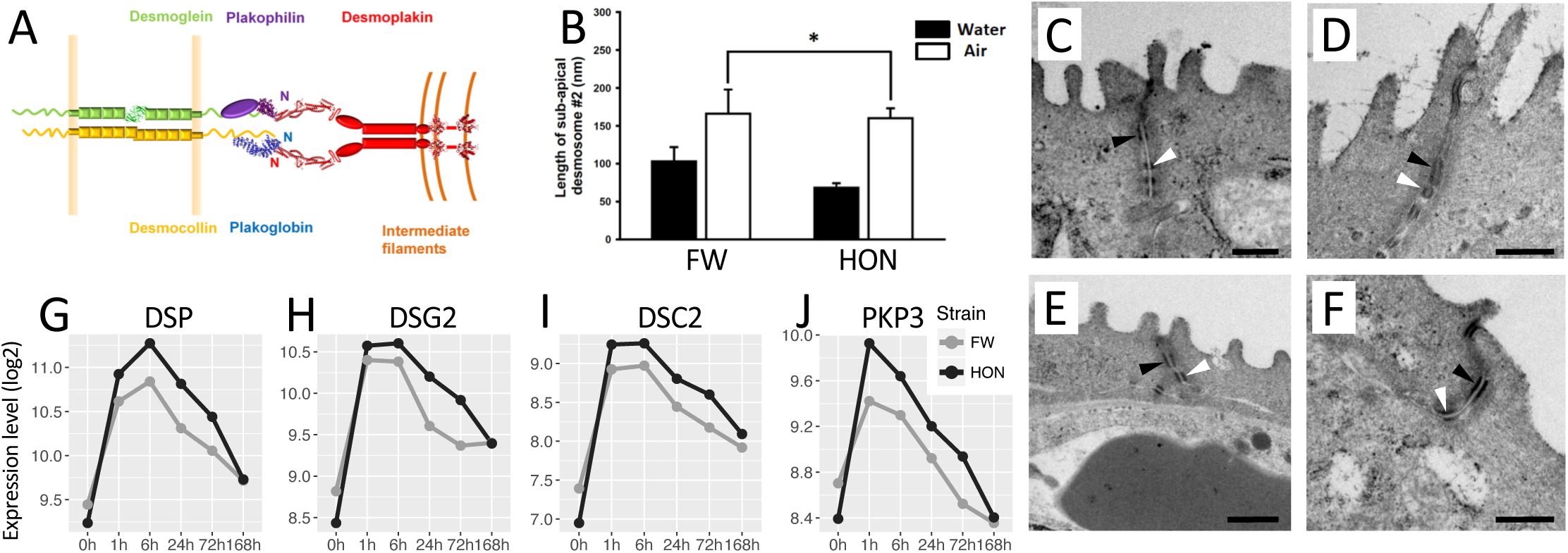
Epidermal desmosome structure and gene expression changes during aerial acclimation. A) proteins that constitute the desmosome (from Al-Jassar C, Bikker H, Overduin M, Chidgey M (2013) Mechanistic basis of desmosome-targeted diseases. J Mol Biol 425(21):4006–4022). B) Length of second desmosome (mm; means ± s.e.m) increases by 7 days in air (open bars) relative to pre-emersion controls (black bars) (*p=0.001). Representative TEM images of skin desmosomes are shown for (C) FW fish at 0 hr, (D) FW fish at 168 hr, (E) HON fish at 0 hr, and (F) HON fish at 168 hr (all scale bars = 500 nm). Desmosome #1 and #2 are indicated by black and white arrowheads, respectively. Log2 expression levels for genes desmoplakin (G), desmoglein-2 (H), desmocollin-2 (I), and plakophilin-3 (J), after time in air (hours) and in pre-emersion controls (0h) for HON (black) and FW (grey) fish.

Molecular pathways and epithelial structures associated with cell-to-cell connections were regulated during emersion. Transcripts for all of the protein subunits that constitute the desmosome were up-regulated by 1 hr post-emersion (Fig. 3), and the GO term *cell-cell adherens junction* and canonical pathway *epithelial adherens junction signaling* are enriched among transcripts up-regulated by 6 hrs post-emersion (Fig. 2A, Table S2). TEM shows deep tight junctions between apical filament-containing cells (keratinocytes) that are exposed to the surrounding environment and numerous desmosomes that ‘stitch’ these and adjacent epidermal cells together in both control and aerially acclimated animals (Fig. 3). Following 7 days of aerial acclimation, morphometric analysis shows that the length of the second sub-apical desmosome (#2) increases in both HON and FW animals (Fig. 3). Desmosomes serve as the intercellular glue that is required for epithelial integrity (27), and desmosome proteins are evolving by directional selection in *K. marmoratus* (see below). This allows us to conclude that connections between keratinocytes that interface with the environment are, at least in part, regulated through desmosomes and that this is likely to be adaptively important for mechanical reinforcement of the skin during emersion.

Structural constituents of the extracellular matrix (ECM) are regulated during aerial acclimation. GO terms *collagen* and *extracellular matrix organization* were enriched among transcripts down-regulated between 1 and 7 days post-emersion (Fig. 2A, Table S6), including the down-regulation of at least 23 collagen transcripts. This suggests that reduced collagen synthesis is an important component of skin remodeling during emersion, which is consistent with our observed reduction in collagen staining in skins sections from both strains by 7 days post-emersion (p<0.001, Fig. 4). TEM observations indicate that collagen is prominent in at least four regions of *K. marmoratus* skin: the stratum compactum in the lower reaches of the dermis, the scales, the dermis/epidermis interface, and epidermal and sub-epidermal vasculature (Fig. 4). Though additional studies will be necessary to definitively identify skin regions where collagen decreases in emersed fish, gene expression data implicate regulation of structures at the dermal/epidermal interface during emersion. Integrins are structural and signaling molecules important for focal adhesion complexes that anchor epidermal cells to the ECM (including collagen) at the dermal/epidermal boundary. The KEGG pathway *focal adhesion* is enriched for genes up-regulated by 6 hrs post-emersion (Fig. 2A, Table S2), including transcripts for integrin subunits and integrin ligands (Figs. S2, S3). We notice that collagen in the sub-epidermal region of *K. marmoratus* skin exhibited an unusual structural arrangement relative to that reported for other fishes, as it appears in loosely ordered bundles (Fig. S6), compared to the highly ordered and deep plywood-like lamellae localized to the upper reaches of the dermis in other fishes (e.g., (28–30)). We conclude that emersion quickly activates pathways that regulate junctional structures between epithelial cells and the ECM, and that atypical sub-epidermal architecture, epithelial remodeling, and morphogenesis are crucial elements underpinning rapid aerial acclimation in *K. marmoratus*.

**Figure 4.**
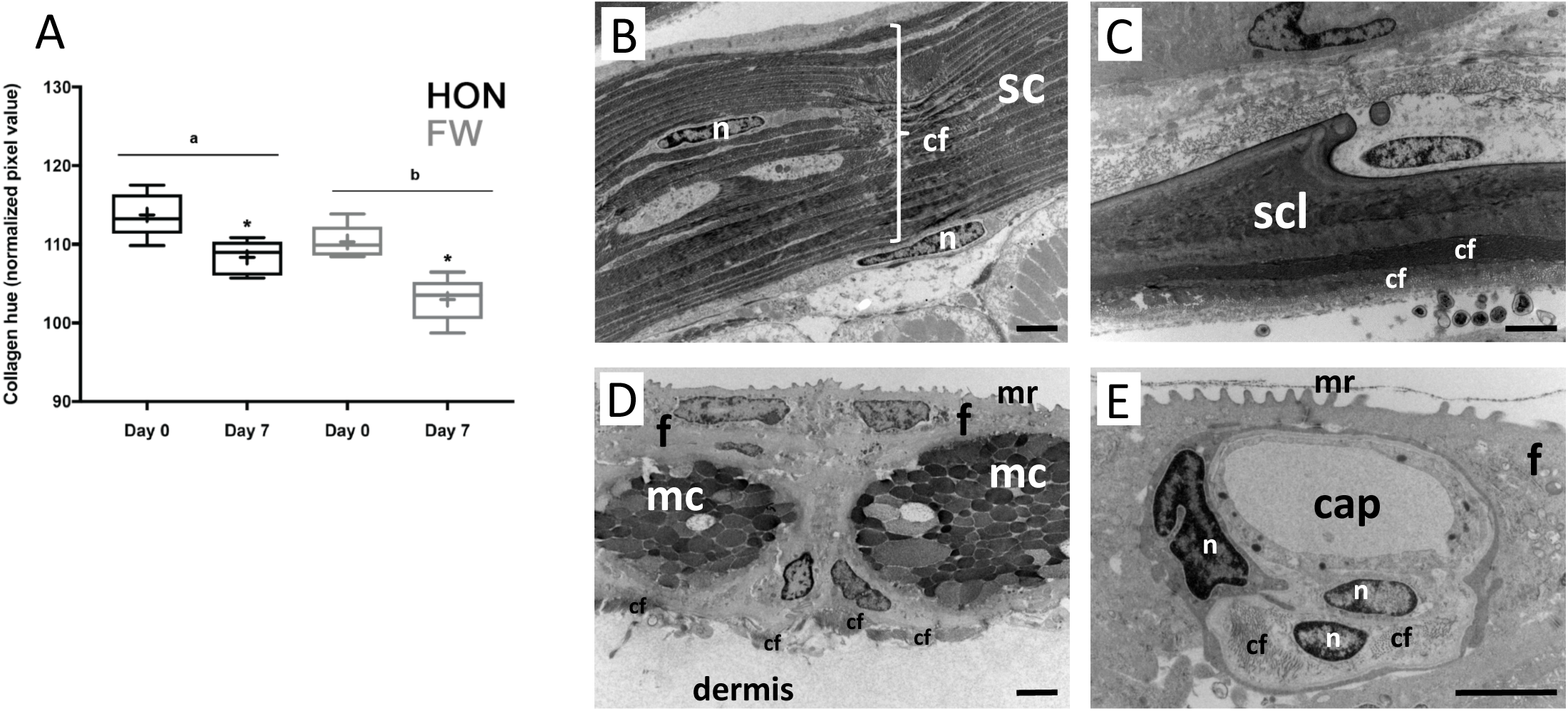
Observations of skin collagen abundance and sites of conspicuous collagen presence. A) skin collagen staining indicates lower levels in FW fish (gray boxes) than HON fish (black boxes) (letters, p=0.01) and a decrease in abundance during emersion in both strains (* p<0.01). Box plots represent 25% and 75% percentiles (+, mean; ___ median); the upper and lower whiskers extend to the minimum and maximum values. TEM images show conspicuous collagen abundance in four regions of *K. marmoratus* skin, including B) the stratum compactum, C) scales, D) collagen fibers loosely dispersed/arranged at the base of the epidermis in the upper reaches of the dermis (i.e. sub-epidermal at the epidermis/dermis interface but located within the dermal region) and E) collagen fibers associated with integumentary blood vessels. sc = stratum compactum; scl = scale; cf = collagen fibers; cap = blood capillary; f = filament-containing cell; mc = mucous cell; mr = microridge; n = nucleus. All scale bars = 2 µm.

As vertebrate life emerged onto land, a key challenge was to prevent water loss; in amniotes, the stratum corneum (SC) forms the outermost layer of the epidermis and functions as a barrier to protect from dehydration. Evolution of this barrier function of the SC is considered crucial for the terrestrial radiation of the amniotes (31). Though fishes lack many of the key proteins that constitute the mature SC in amniotes (e.g., loricrin, filaggrin, involucrin) (32) transcriptional data provide evidence that the molecular machinery that supports formation of the SC in amniotes is activated during emersion acclimation in *K. marmoratus* skin. In mammalian epidermis, the transglutaminases (in particular TGM1 and TGM5) govern the process of protein cross-linking that produces the SC. These proteins have been found in fish, are expressed in epithelial tissues, and TGM1 appears to retain the ability to form cross-linked protein structures (33). In *K. marmoratus* skin, TGM1 (which contains the characteristic N-terminal cysteine cluster) and TGM5 are significantly up-regulated within 1 hr of aerial exposure (Fig. S7). The other key proteins that regulate this pathway are also up-regulated by 1 and 6 hrs post-emersion (e.g., EVPL, PPL, KAZN; Fig. S7). Furthermore, the sphingolipid metabolism pathway, which produces lipids integral to barrier function of the SC in amniotes (34), is up-regulated between 1 and 7 days post-emersion (Fig. 2A, Table S3). TEM images provide no evidence for the formation of a stratum corneum in 7-day acclimated *K. marmoratus*, so the physiological manifestations of these SC-associated molecular responses merit further inquiry.

### Ammonia detoxification

In water, metabolic waste in the form of toxic ammonia is efficiently excreted across the gills. However, alternate mechanisms are required to avoid ammonia poisoning in the absence of gill-water contact, such as for species occupying terrestrial niches. These mechanisms could include ammonia excretion across the skin, either by acid trapping or volatilization, or metabolic conversion to less toxic forms such as urea or glutamine (35).

Models for ammonia excretion in aquatic species were updated with the discovery of the ammonia-transporting properties of Rhesus (Rh) glycoproteins (36). In *K. marmoratus*, the three Rh glycoproteins Rhag, Rhbg, Rhcg show increased expression by 6 to 24 hrs post-emersion (p=0.003, 0.03, 0.0004) (Fig. S8). Rh glycoprotein expression responses are correlated with those of V-type H^+^-ATPase subunits (Fig. S8), where combined these proteins could function as an ammonium pump, consistent with an acid-trapping model for ammonia excretion proposed for fish gills (36). Rh glycoproteins are localized to the apical crypt of epidermal ionocytes in *K. marmoratus* (37); we posit that the deep apical crypts of ionocytes that we observed in air-exposed fish (Fig. 5A) may be important for creating a suitable microenvironment for the acid trapping mechanism. Careful regulation of skin surface pH is likely important to balance acid trapping of NH_3_ and volatilization of gaseous NH_3_ in air (38), and this is achieved through H^+^-ATPase and Na^+^/H^+^ exchangers (NHE) (39). Expression of NHE2 is over 50-fold higher in HON fish than in FW fish (Fig. S9). Since inhibition of NHE2 expression affects skin pH in air and disrupts ammonia excretion across the skin (39), this difference in NHE2 expression between HON and FW fish could reflect differences in the ability to excrete ammonia across the skin in air and thereby contribute to differences in terrestrial acclimation abilities between strains.

**Figure 5.**
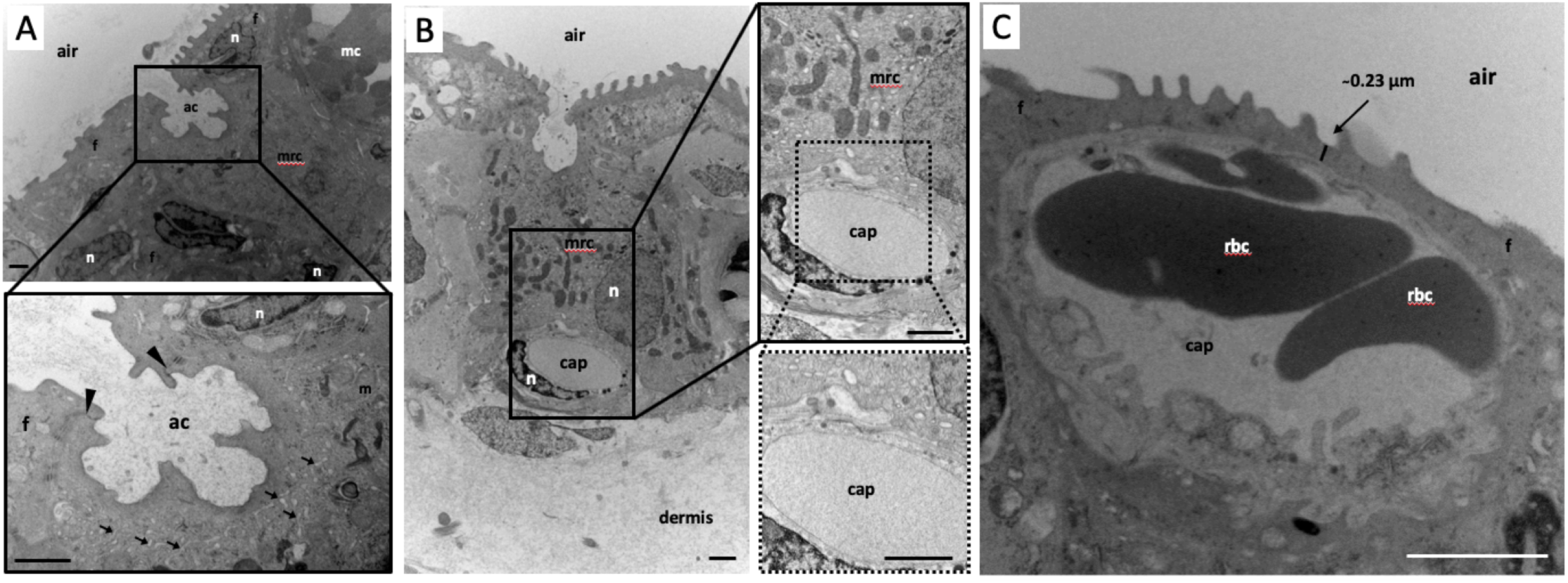
TEM observations of epidermal ionocytes in air-exposed fish and skin blood vessels. Deep apical crypts are observed in air-exposed ionocytes of (A) FW and (B) HON fish. In both FW and HON fish, a dense intracellular tubular network is indicated by black arrows, and deep tight junctions (A, black arrowheads) link ionocytes to adjacent epidermal cells. Panel (B) shows a blood capillary residing at the base of an ionocyte while panel (C) shows a blood capillary extremely close to the apical surface of the epidermis, resulting in diffusion distances as low as 0.23 µm. Scale bars in (A) and (B) = 1 µm and in (C) = 2 µm. ac = apical crypt; cap = capillary; f = filament-containing cell; m = mitochondrion; mc = mucous cell; mrc = epidermal ionocyte; n = nucleus.

In addition to excretion, ammonia toxicity may be ameliorated through metabolism to less-toxic substances. For example, glutamine synthetase (GLUL) converts ammonia to non-toxic glutamine (40), thereby providing a crucial ammonia detoxifying system. In *K. marmoratus*, expression of GLUL is significantly increased at 1 hr post-emersion and stays high throughout their time in air (Fig. S10). These data are consistent with Rh glycoprotein-mediated ammonia excretion, coupled with metabolic detoxification through glutamine synthetase, which enables regulation of ammonia across the skin during acclimation to the terrestrial environment.

### Water/ion transport

*K. marmoratus* are able to effectively counteract desiccation while out of water; after several days in air they increase water influx across the skin, whereas water efflux does not change (41), and fish maintain whole-body water levels (42). Enhanced water uptake may involve aquaporins or changes to paracellular tight junctions (43, 44), and upon emersion we observed rapid up-regulation of aquaporin 3, as well as claudin 10 and occludin tight junction transcripts (Figs. S11, S5).

In air-exposed fish, TEM revealed two types of sub-apical tight junctions in the epidermis. The most common were deep tight junctions, which link most cells in the epidermis. Deep tight junctions are comparatively long (∼ 200 nm) and appear to either completely occlude the intercellular space (Fig. S12A) or reduce it so that it appears as a single membranous “line”. In contrast, a second type of tight junction observed were shallow (“leaky”) tight junctions, which linked ionocytes and adjacent interdigitating cells in an arrangement that is similar to the relationship between seawater fish gill ionocytes and interdigitating accessory cells. In *K. marmoratus* skin, shallow tight junctions directly interface with the environment within ionocyte apical crypts (Fig. S12B,C). Shallow tight junctions were frequently observed in TEM images from HON animals acclimated to air, but none were observed in air-acclimated FW animals, despite the presence of apical crypts in these animals. Indeed, this latter observation is particularly novel because in the gill epithelium of fishes, ionocyte apical crypts are a hallmark of seawater but not FW gill architecture. This is because in the seawater fish gill epithelium, ionocyte apical crypts establish a microenvironment within which paracellular movement of Na^+^ across leaky junctions can occur, down an electrochemical gradient established by transcellular Cl^-^ secretion (45). Therefore the enigmatic presence of deep apical crypts without “leaky” junctions in ionocytes of air-exposed FW fish most likely reflects the need to create a microenvironment for solute transport, not for Na^+^ secretion but possibly for NH_3_ trapping. In line with these observations, HON and FW animals differed in their emersion-induced expression of key components of fish ionocytes, including Na^+^/K^+^-ATPase and CFTR. Apical CFTR is acknowledged to establish the electrochemical gradient that allows Na^+^ transport through leaky tight junctions, and claudin-10 proteins are proposed to selectively facilitate Na^+^ movement (46, 47). Notably, claudin-10 mRNA also exhibits strain and emersion characteristics that are consistent with CFTR (e.g. both CFTR and claudin 10 are more abundant in HON versus FW animals). Therefore, our observations suggest that Na^+^ regulation through “leaky” tight junctions that reside within apical crypts of HON epithelial ionocytes is important for aerial acclimation in this strain. Indeed, many genes that are typically expressed in fish ionocytes, and that facilitate water and salt homeostasis, were expressed differently between HON and FW fish in response to emersion (see *Different responses to emersion between HON and FW fish* section below). We conclude that effective regulation of water and ions enables aerial acclimation, and that variable osmoregulatory strategies may be linked to differences in resilience to emersion between HON and FW animals.

### Respiration

If fish skin is to act like a gill on land, then oxygen transport through this organ should be enhanced during aerial acclimation. In this regard, a morphological hallmark of cutaneous respiration in vertebrates is augmented vascularization of the skin, and in some cases the presence of blood vessels in the upper reaches of the epidermis which reduces the blood to water diffusion distance and enhances epithelial gas transport in air (48). In this as well as previous studies, epidermal blood vessels have been observed in *K. marmoratus* (49). However, our TEM images show that by 7 days post-emersion the diffusion distance from air to blood in epithelial capillaries is as short as 0.23 μm (Fig. 5C); to our knowledge, this is the shortest cutaneous diffusion distance yet measured in a fish. By comparison, diffusion distance across the skin of other amphibious fishes, as well as the gill epithelium of fishes in general, is at least one order of magnitude greater (50, 51). Fish respiratory organs that have been reported to match the blood to water diffusion distance of *K. marmoratus* skin are those of the suprabranchial chamber and labyrinthine organ of the climbing perch (50).

Cutaneous angiogenesis was elevated within one day of air exposure in *K. marmoratus*, accompanied by rapidly elevated expression of key angiogenesis regulating genes including VEGF, ANGPT, EFNA, and PECAM1 (8). VEGF plays a fundamental role in angiogenesis (52, 53) and VEGFA itself was up-regulated by 1 hr post-emersion (8). Genes involved in the KEGG pathway *VEGF signaling* were enriched among the genes significantly up-regulated by 6 hrs post-emersion (Table S2), and *angiogenesis* and *vasculogenesis* are BioFunctions that were significantly enriched (p=3.69E-8, 4.65E-9, respectively, IPA analysis) among the genes up-regulated by 24 hrs post-emersion (Fig 2A). These include transcription factors XBP1, NR4A1, and SRF (Fig. S13) that regulate VEGF-dependent angiogenesis (54–56). Furthermore, bradykinin receptor promotes angiogenesis through nitric oxide signaling (57), and we find parallel up-regulation of BDKRB2 transcripts and NOS1 by 6 hrs post-emersion (Fig. S13). We conclude that angiogenesis, coupled with blood vessels in very close proximity to the apical surface (Fig. 5C), is important for shifting respiratory functions from the gill to the skin during aerial acclimation.

In addition to blood vessels being located in the upper reaches of *K. marmoratus* skin (Fig. 5C) presumably for the purpose of enhancing respiration, TEM observations also revealed the presence of blood vessels adjacent to ionocytes. When ionocytes were observed in the epidermis of air-exposed *K. marmoratus* skin, blood vessels were usually lying adjacent and in direct contact with them (Fig 5B). Blood vessels were typically located at the base of the ionocyte at the epidermal-dermal border (but within the dermis). This arrangement essentially establishes the ionocyte as a blood-to-water bridge, across which solutes could be efficiently transported. Functionally, this is analogous to the gill epithelium where an exposed ionocyte interfaces with water at the apical surface and the blood of the gill vasculature at the basolateral surface (58). Therefore, from an architectural standpoint, in addition to observing a reorganization of cutaneous vasculature for respiratory purposes, we also observe a reorganization of integument vasculature to presumably enhance solute transport properties. The highly organized spatial arrangement of blood vessels, both very close to the skin surface and associated with ionocytes, is likely to be functionally important to air-exposed *K. marmoratus*.

### Different responses to emersion between HON and FW fish

Considering 7-day survival data, the HON fish were more resilient to emersion than FW fish, and resilience correlated with reduced metabolic rate during acclimation (Fig. 1, Turko et al. in review). Though resilience to emersion is likely supported by physiological responses in multiple organs, we examined genes that were differentially expressed between strains during emersion to offer insight into the mechanisms that may contribute to this variable resilience in the skin. Among the genes that were differentially expressed between strains during aerial acclimation (strain*time interaction, p<0.05), we observe three clusters of co-expressed genes (Fig. 2B); Set 1: these genes show a transient spike in expression at 1 hour for HON fish, but gradual down-regulation then recovery in FW fish. Set 1 is significantly enriched for GO terms *cell cycle* (p=8.6E-56) (and other related GO terms: Table S7). Set 2: these genes show gradual up-regulation only in FW fish with little temporal response in HON fish. Set 2 is enriched for GO terms *proteasome* (p=5.3E-18) and *lysosome* (p=9.4E-4) (Table S8). Set 3: these genes show gradual down-regulation only in FW fish with little temporal response in HON fish. Set 3 is enriched for GO terms *translational initiation* (p=8.7E-6), *cell junction* (p=7.7E-5), and *sodium transport* (p=3.1E-2) (Table S9). In addition to the sodium transport genes (including Na^+^/K^+^-ATPase, Na^+^-bicarbonate cotransporter, Na^+^/H^+^ exchanger) we detect a number of other genes known to be involved in osmoregulation in fish gills and differentially expressed between strains, including V-type H^+^ ATPase, CFTR, carbonic anhydrase, claudins, and aquaporin (Fig. S14). Set 3 is also enriched for the KEGG pathway *ribosome*, which is a core part of the transcriptional environmental stress response in yeast (59).

Structural differences of the skin between FW and HON fish, because of differences in acclimation or adaptation to fresh water, may also contribute to their differing acclimation abilities. For example, ionocyte-associated shallow “leaky” tight junctions were frequently observed in TEM images from HON animals acclimated to air, but none were observed in air-acclimated FW animals; this is typical of gill ionocytes acclimated to salty water. Since shallow tight junctions regulate sodium transport, this is consistent with divergent patterns of expression of ionoregulatory genes outlined above.

We propose the following to explain differences in resilience to emersion between HON and FW fish: immediate differences in cell cycle regulatory response may indicate differences between HON and FW animals in their ability to sense and respond to cellular stress incurred upon emersion (set 1). Cellular damage may have accumulated in the less-resilient FW strain, such that proteolytic activity is up-regulated primarily in that strain (set 2). Differences between HON and FW animals in their regulation of structural and functional components of the osmoregulatory apparatus in ionocytes (set 3) contributes to differences in resilience to the ionoregulatory challenges posed by emersion. In particular, genes that code for proteins that are directly linked to sodium uptake (e.g., NHE, NBC and accessory protein CA, NKA; (60)) showed a downward or erratic trend in our FW strain. If the skin is solely responsible for sodium uptake in air-exposed fish as we presume, then a lack of a clear concerted induction of the NHE/NBC pathway may indicate that sodium uptake was limited in FW fish which may have affected survival. Though plasma sodium is difficult to measure in very small fish, additional studies are merited to test this hypothesis. We conclude that osmoregulatory differences from acclimation and/or adaptation to fresh water constrains the ability of *K. marmoratus* to adjust to emersion.

### Protein evolution

Our initial test for positive selection in *K. marmoratus* protein sequences included 1,073 single-copy orthologous gene loci in all nine fish species included (Table S12, Fig. S15), and revealed 24 genes showing positive selection in the *K. marmoratus* lineage only (Table S13, Fig. S15). An extended analysis expanded the query set to include orthologs present in all species except one of either *D. rerio*, *T. rubripes*, or *G. aculeatus*; this included 461 further genes for analysis (Table S14), and revealed 16 more gene loci evolving by positive selection in the *K. marmoratus* lineage (Table S15, Fig. S15). Because of the constraints for confidently identifying single-copy orthologs across multiple taxa, this analysis queried only ∼4% of the genes in the *K. marmoratus* genome (Table S16). Therefore, we likely did not detect many of the genes evolving by positive selection within this unusual species. For example, Rhesus glycoproteins appear to be evolving by positive selection in amphibious mudskippers (61), but since these proteins were not included in our query set we cannot infer whether they similarly show accelerated evolution in amphibious *K. marmoratus*. However, we predicted that some of the genes evolving by positive selection in the *K. marmoratus* lineage would contribute to the unique terrestrial abilities of this species, such as those involved in the skin functions implicated by gene expression, physiological assays, and structural imaging reported here.

We detected positive selection in genes involved in skin morphology and morphogenesis, including two key genes that contribute to desmosome structure (desmoglein and plakophilin) and in transcription factor Kruppel-like factor 9 (KLF9) that regulates skin morphogenesis (62). Desmosomes share a conserved role in maintaining epidermal integrity to mechanical stress in other vertebrates including amphibians (63), and are subject to selection during re-invasion of aquatic environments in mammals (64). Furthermore, human diseases of the heart and skin are associated with mutations in desmosome structure and expression, and with desmosome genotype-by-environment interactions (65). Given that desmosome proteins are evolving by directional selection in the amphibious *K. marmoratus*, they may prove a valuable model for exploring the eco-physiological importance of desmosome regulation, structure, and function. Positive selection was also detected in heart- and neural crest derivatives-expressed protein 1 (HAND1) which is required for angiogenesis, and regulates the expression of many angiogenic pathways including VEGF, angiopoietin, and ephrin signaling (66). Pathways related to skin remodeling, desmosome function, and angiogenesis are also among those showing patterns of differential regulation during terrestrial acclimation (Fig. 2A). We conclude that proteins evolving by positive selection within the *K. marmoratus* lineage are adaptive for supporting skin plasticity that enables the amphibious lifestyle of this species.

### Summary

Amphibious abilities have evolved in at least 33 families of extant fish (1, 67, 68), many species of amphibians transition between aquatic and terrestrial forms during metamorphosis, and three lineages of terrestrial mammals have transitioned back to fully aquatic forms (cetaceans, pinnipeds, and sirens). This diversity provides ample opportunity to study the conserved and divergent mechanisms that support transitions between aquatic and terrestrial physiologies that are so important in the history of vertebrate diversification. So far, few studies have detailed the mechanisms that underlie skin plasticity and evolution during emersion. Our integrated molecular, structural, and evolutionary analyses reveal the importance of regulation of cell-cell adhesion structures, regulation of molecular pathways that govern skin morphogenesis, and evolution of desmosome proteins to mechanically reinforce and buttress barrier functions of this tissue. The challenge of supporting respiratory and ion exchange functions is associated with ionocyte structures and rapid regulation of ionoregulatory pathways, angiogenesis, evolution of angiogenesis-regulating proteins, and an unusual blood vessel architecture that links very short air-blood diffusion distance with delivery to energetically demanding ionocytes. Studies in marine mammals and amphibious gobies provide evidence of parallel importance of skin morphogenesis (64) and ion transport mechanisms (61), respectively, in aquatic-terrestrial transitions. Future studies could focus on organs in addition to skin, such as eyes, brain, liver, kidney, and muscle, that may contribute to terrestrial acclimation.

We found that *K. marmoratus* derived from brackish niches were more resilient to emersion than freshwater types. Considering that Devonian tetrapods were likely derived from marine or brackish lineages (e.g., (69–71)), and that brackish and terrestrial environments share challenges posed by dehydration, perhaps salty aquatic environments favor osmoregulatory physiologies that are more easily co-opted for terrestriality than freshwater environments. Our data are consistent with this hypothesis and suggest that physiological differences in ionoregulation are mechanistically associated with terrestrial acclimation abilities.

## Acknowledgements & Funding

We thank Dr. Lisa Johnson for bioinformatics advice and training, Jennifer Roach for assistance with RNA isolation and RNA-seq library preparation, Norbert Grundmann for assistance in using the PALMA cluster Muenster, Chun Chih Chen for assistance with morphometric work, and Drs. Nick Bernier, Beren Robinson, and Andy Turko for advice on experimental design and metabolic rate measurements. This work was supported in part by grants from the U.S. National Science Foundation (OCE-1314567 and DEB-1265282 to A.W.), the U.S. National Institutes of Environmental Health Sciences (1R01ES021934-01 to A.W.), the Deutsche Forschungsgemeinschaft (SCHM1469/10-1 to J.S.), and the Natural Sciences and Engineering Research Council of Canada (RGPIN-2018-04218 to P.W., RGPIN 2014-04073 to S.P.K.). T.B. was supported by an Ontario Graduate Scholarship.

## Figure legends

**Figure S1.**
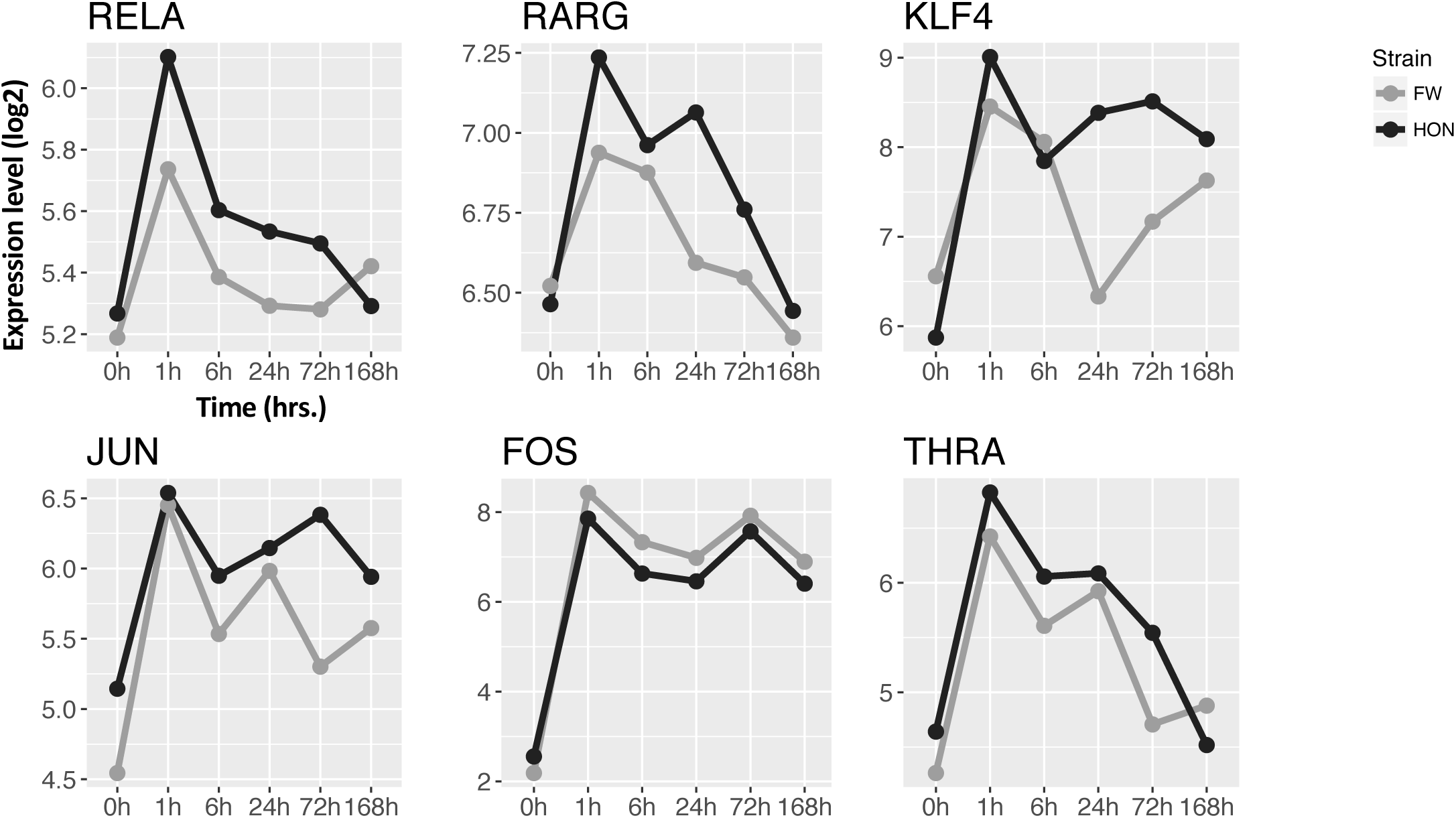
Log2 expression levels for skin morphogenesis transcription factor genes, including transcription factor p65 (RELA), retinoic acid receptor gamma (RARG), krueppel-like factor 4 (KLF4), transcription factor AP-1 (JUN and FOS), and thyroid hormone receptor alpha (THRA), after time in air (hours) and in pre-emersion controls (0h) for HON (black) and FW (grey) fish.

**Figure S2.**
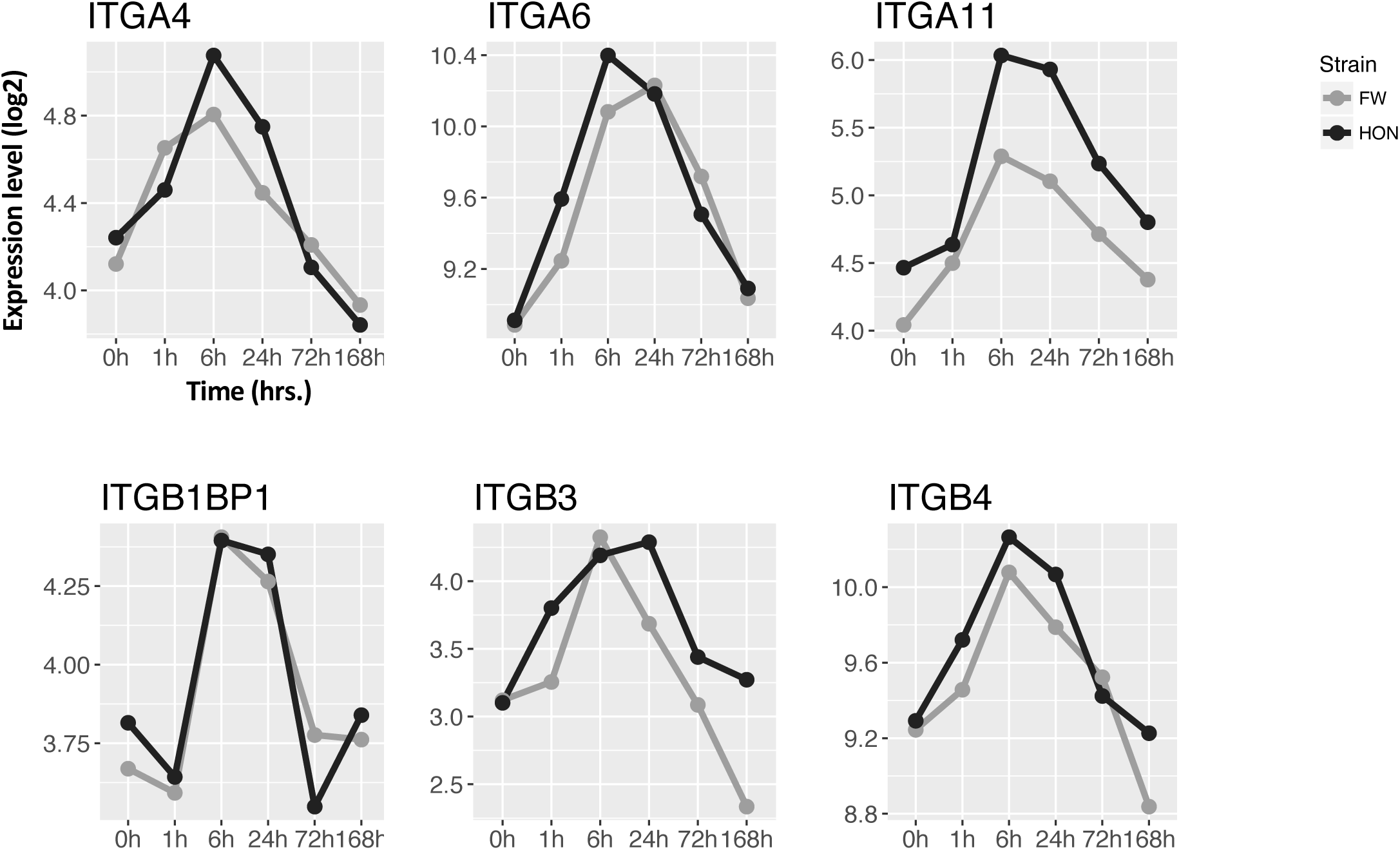
Log2 expression levels for integrin subunit genes after time in air (hours) and in pre-emersion controls (0h) for HON (black) and FW (grey) fish.

**Figure S3.**
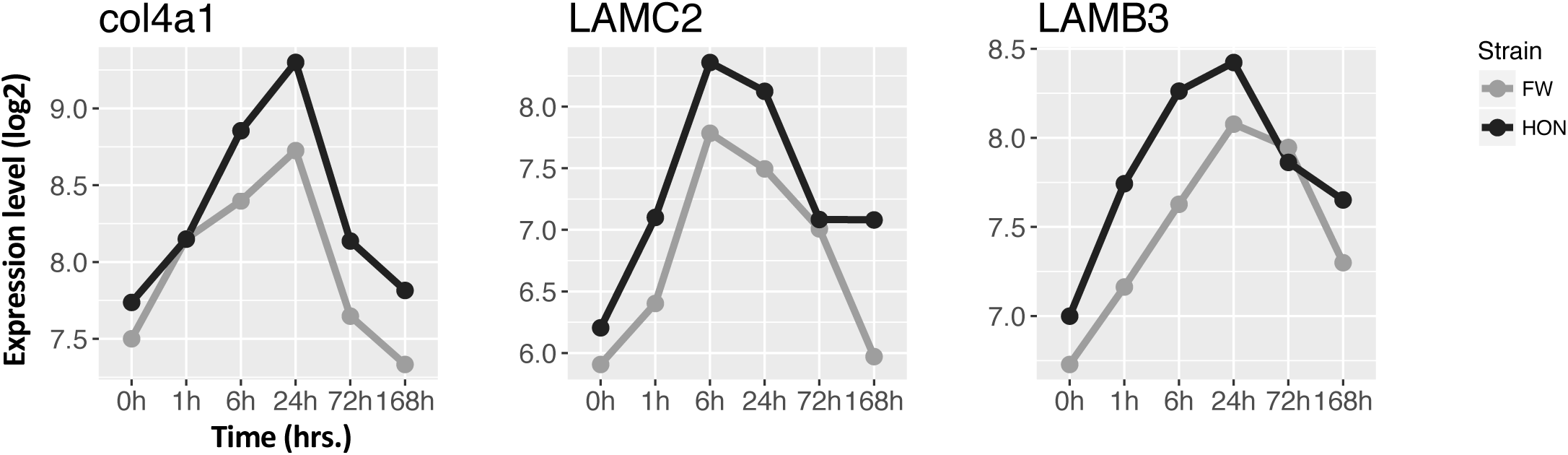
Log2 expression levels for integrin ligand genes, including collagen alpha-1(IV) chain (col4A1), laminin subunit gamma-2 (LAMC2), and laminin subunit beta-3 (LAMB3), after time in air (hours) and in pre-emersion controls (0h) for HON (black) and FW (grey) fish.

**Figure S4.**
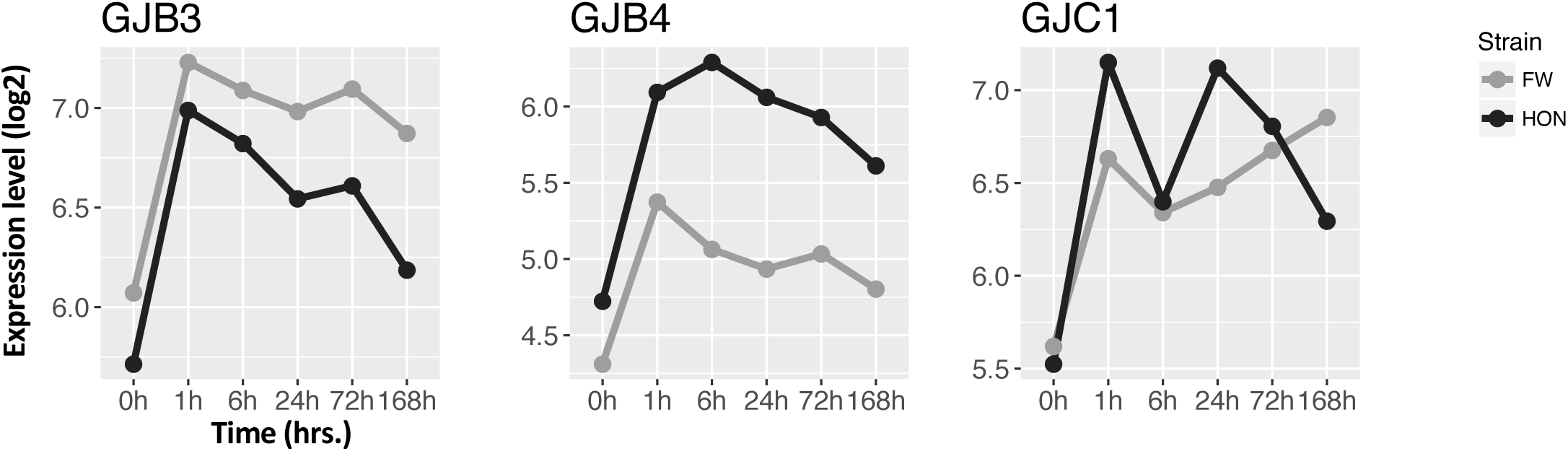
Log2 expression levels for gap junction genes after time in air (hours) and in pre-emersion controls (0h) for HON (black) and FW (grey) fish.

**Figure S5.**
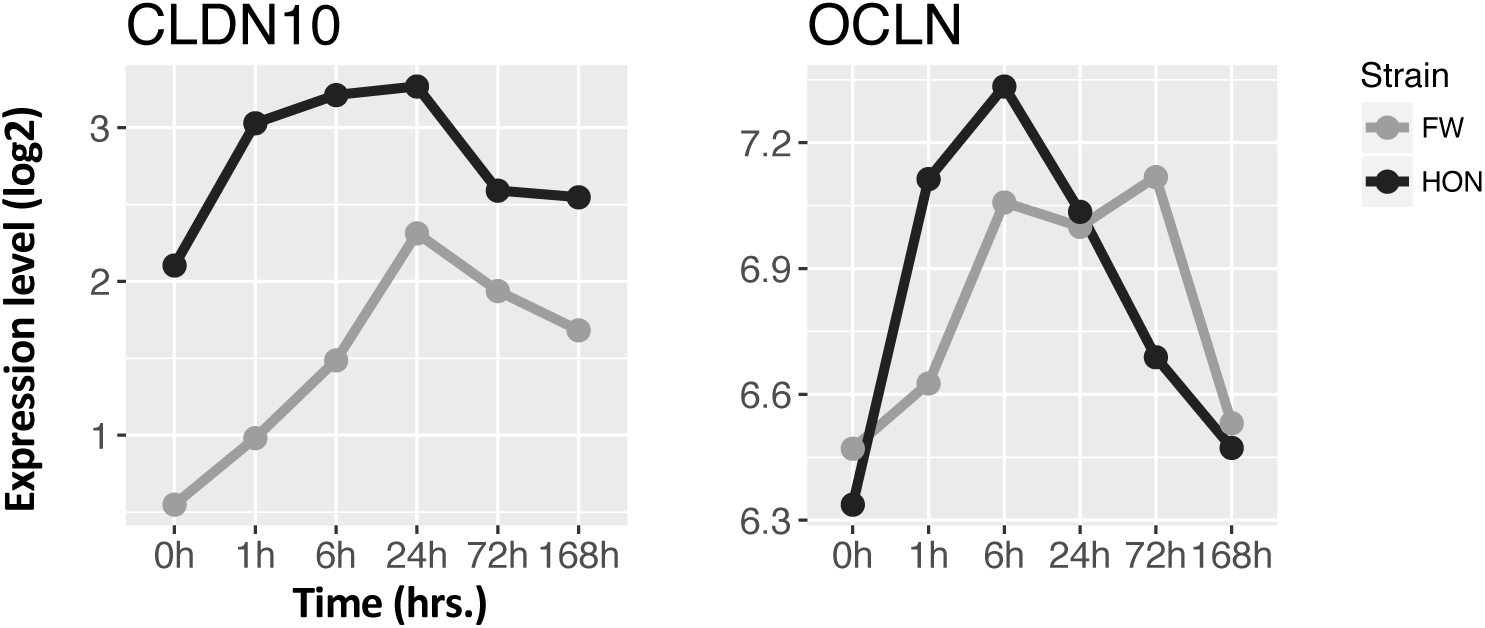
Log2 expression levels for tight junction genes, including claudin-10 (CLDN10) and occludin (OCLN), after time in air (hours) and in pre-emersion controls (0h) for HON (black) and FW (grey) fish.

**Figure S6.**
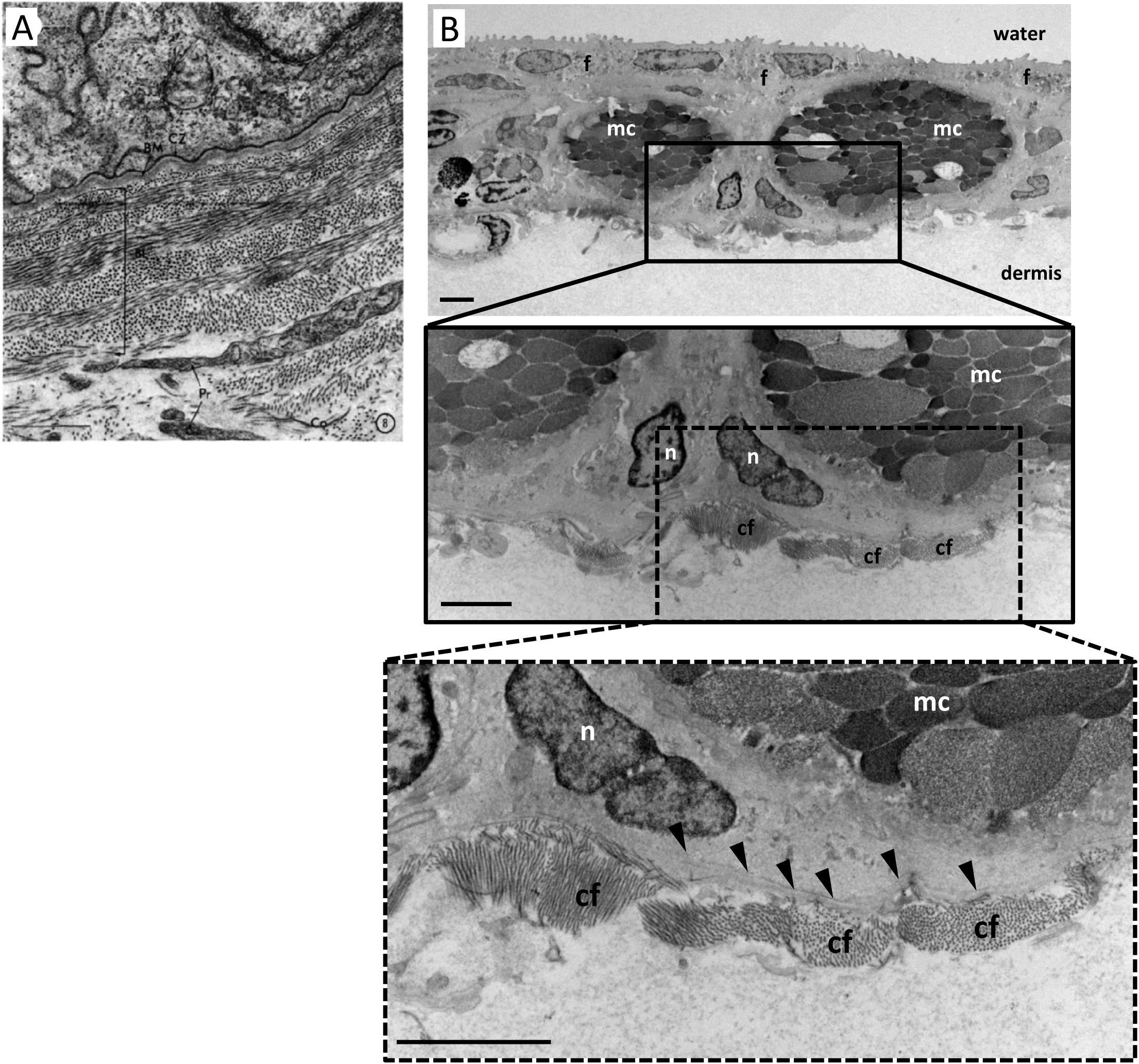
TEM images contrast the highly-structured organization of collagen at the dermal/epidermal boundary of a non-amphibious Cyprinodontiforme, *Fundulus heteroclitus* (A), with the loosely bundled organization of collagen at the dermal/epidermal boundary in *K. marmoratus* skin (B). Scale bars in panel (B) and insets = 2 µm. Images in panel (A) and high magnification inset from panel (B) were taken at x 21 500 and x 14 500 respectively. cf = collagen fibers, f = filament-containing cell, mc = mucous cell, n = nucleus, and the basement membrane is indicated with black arrowheads. Image in (A) is taken from Nadol, J. B., J. R. Gibbins, and K. R. Porter. 1969. A reinterpretation of the structure and development of the basement lamella: An ordered array of collagen in fish skin. Developmental Biology 20:304–331.

**Figure S7.**
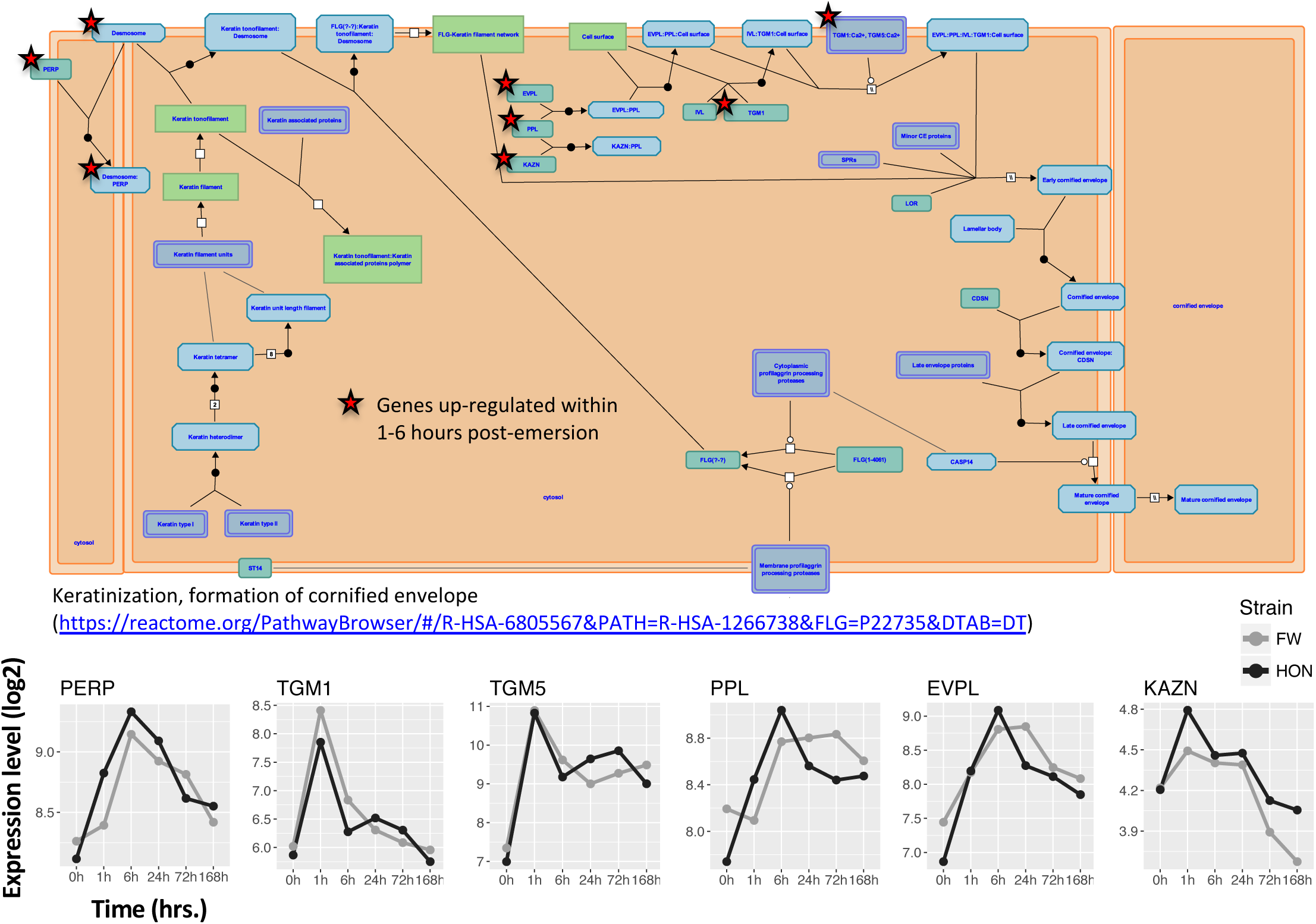
Genes that are part of the pathway that regulates formation of the cornified envelope in terrestrial vertebrates were up-regulated during emersion (red asterisks in pathway map; map is from reactome.org). Line graphs are log2 expression levels for genes, including p53 apoptosis effector related to PMP-22 (PERP), transglutaminase-1 (TGM1), transglutaminase-5 (TGM5), periplakin (PPL), envoplakin (EVPL), and kazrin (KAZN), after time in air (hours) and in pre-emersion controls (0h) for HON (black) and FW (grey) fish.

**Figure S8.**
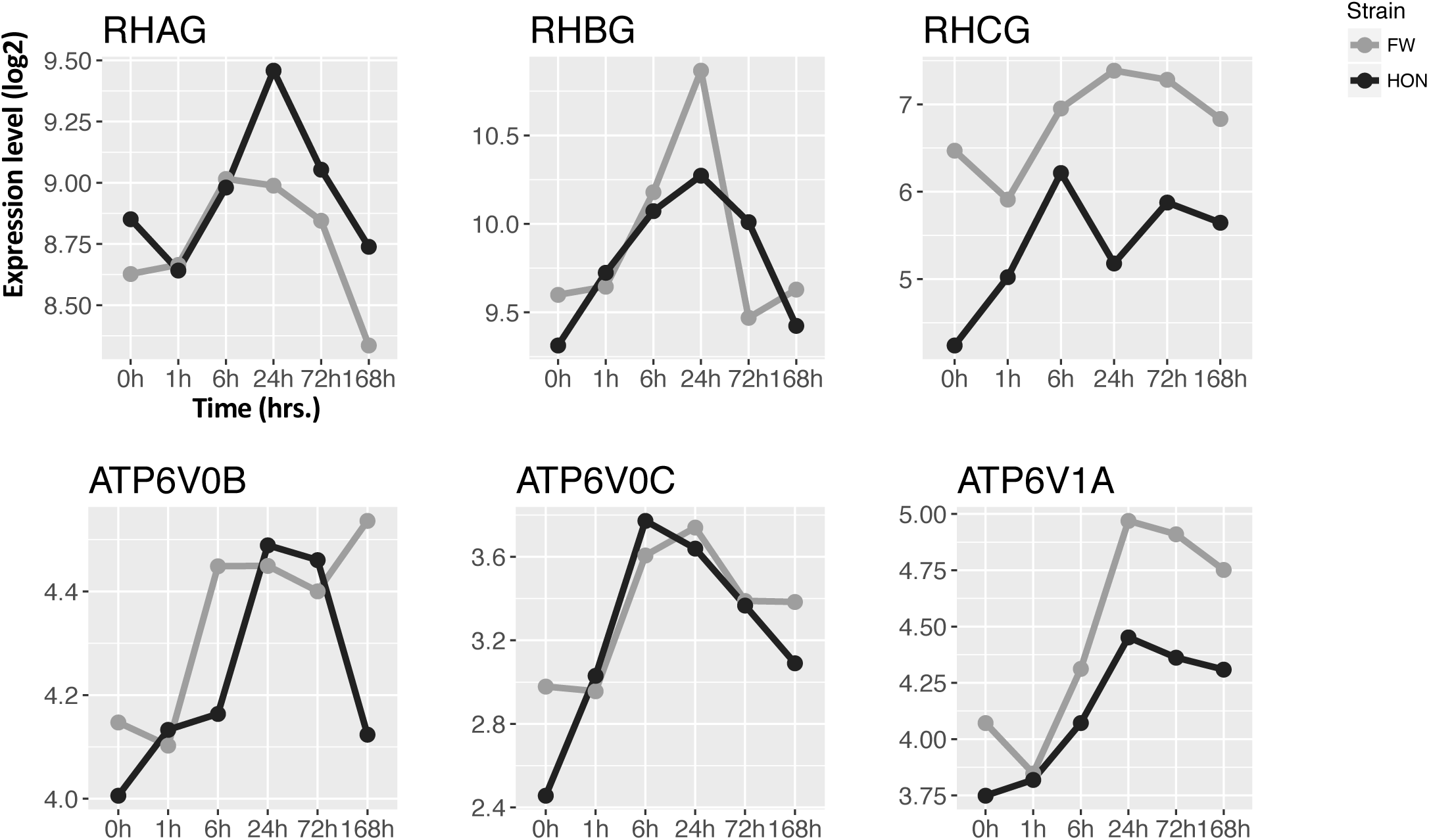
Log2 expression levels for genes, including ammonium transporter Rh type A (RHAG), ammonium transporter Rh type B (RHBG), ammonium transporter Rh type C (RHCG), V-type proton ATPase 21 kDa proteolipid subunit (ATP6V0B), V-type proton ATPase 16 kDa proteolipid subunit (ATP6V0C), and V-type proton ATPase catalytic subunit A (ATP6V1A), after time in air (hours) and in pre-emersion controls (0h) for HON (black) and FW (grey) fish.

**Figure S9.**
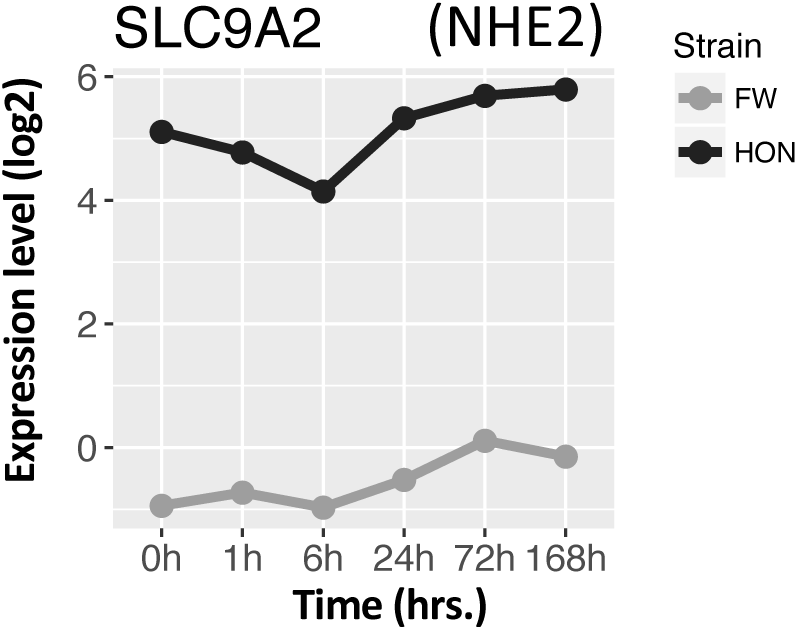
Log2 expression levels for gene sodium/hydrogen exchanger 2 (SLC9A2, or NHE2), after time in air (hours) and in pre-emersion controls (0h) for HON (black) and FW (grey) fish.

**Figure S10.**
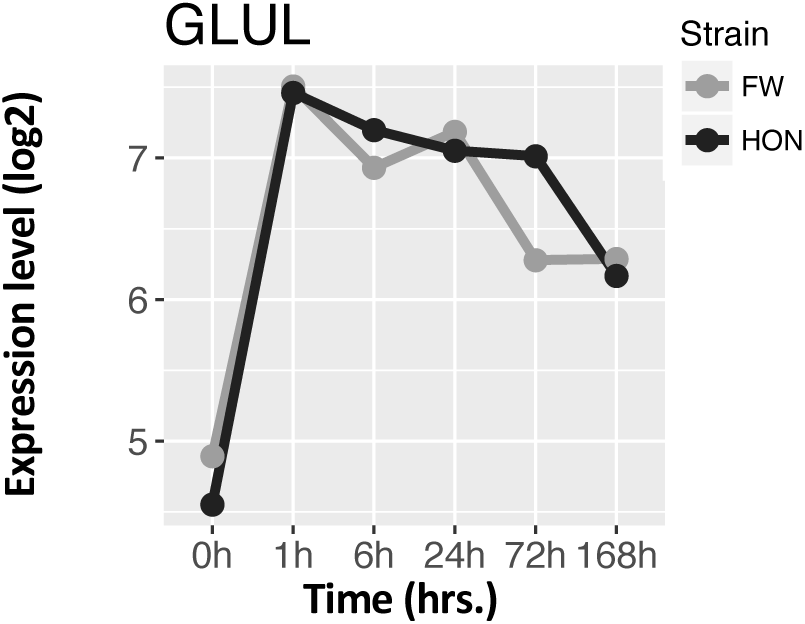
Log2 expression levels for gene glutamine synthetase (GLUL), after time in air (hours) and in pre-emersion controls (0h) for HON (black) and FW (grey) fish.

**Figure S11.**
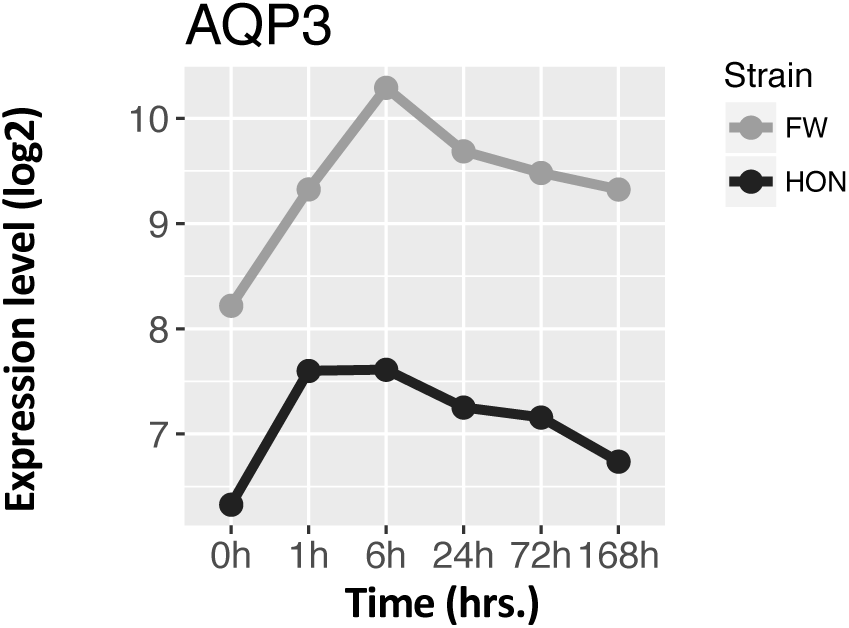
Log2 expression levels for gene aquaporin-3 (AQP3), after time in air (hours) and in pre-emersion controls (0h) for HON (black) and FW (grey) fish.

**Figure S12.**
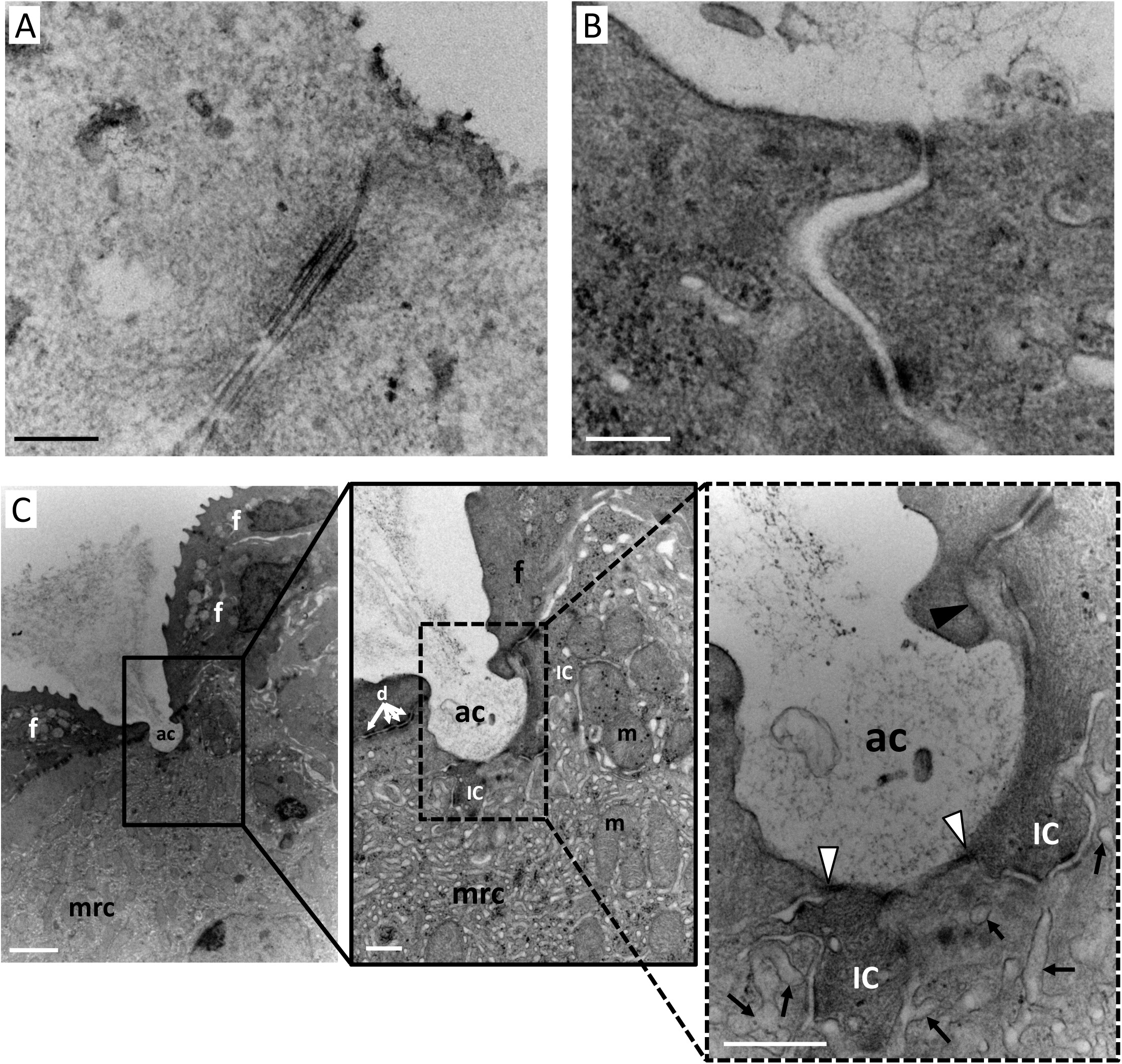
TEM images contrast (A) deep epidermal tight junctions which are located between adjoining keratinocytes or link ionocytes to keratinocytes and (B) shallow “leaky” tight junctions that link ionocytes and adjacent interdigitating cells. Note that deep tight junctions completely occluded the intercellular region while shallow tight junctions exhibit “kissing points” where adjacent cell membranes can be easily distinguished. Panel (C) shows an epidermal ionocyte from an air-exposed HON animal with a deep (simple cup-shaped) apical crypt exposed to the surrounding environment. Shallow tight junctions (white arrowheads) link the ionocyte and an interdigitating cell within the environment of the apical crypt. Numerous shallow tight junctions are observed in HON fish in air, but are not observed in FW fish in air. The dense intracellular tubular network (a tortuous invagination of the basolateral membrane, black arrows) is particularly prominent and it is quite obvious how close to the apical surface of the ionocyte that this structure reaches. Deep tight junctions (black arrowheads) link the ionocyte to keratinocytes/adjacent filament-containing cells. Scale bars = 200 nm for (A) and (B), 2 µm for panel (C) and 500 nm for panel (C) insets. ac = apical crypt; d = desmosome; f = filament-containing cell; ic = interdigitating cell; m = mitochondrion; mrc = epidermal ionocyte; n = nucleus.

**Figure S13.**
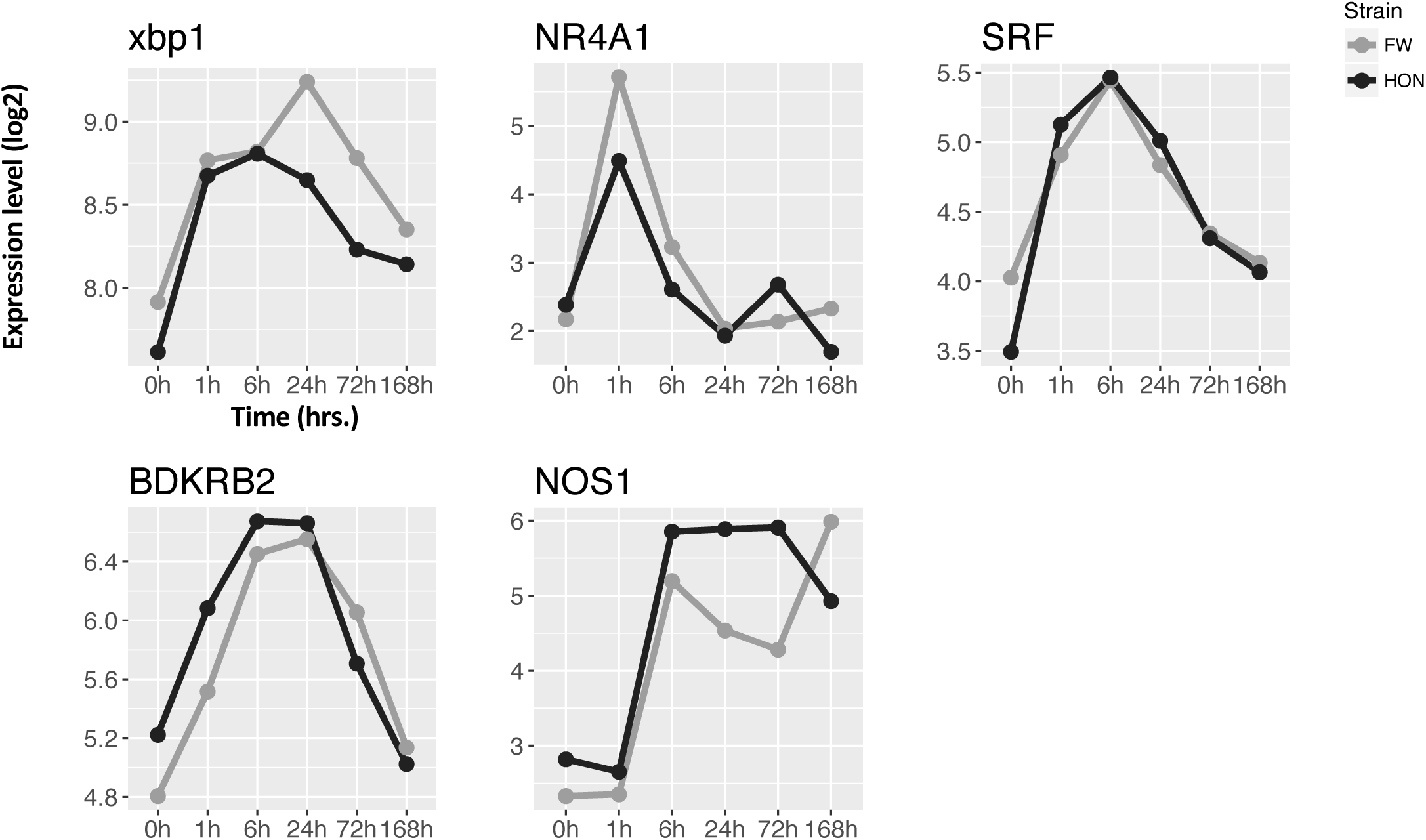
Log2 expression levels for genes encoding transcription factors that regulate VEGF-dependent angiogenesis, including X-box-binding protein 1 (xbp1), nuclear receptor subfamily 4 group A member 1 (NR4A1), and serum response factor (SRF), plus genes B2 bradykinin receptor (BDKRB2) and neuronal nitric oxide synthase (NOS1), after time in air (hours) and in pre-emersion controls (0h) for HON (black) and FW (grey) fish.

**Figure S14.**
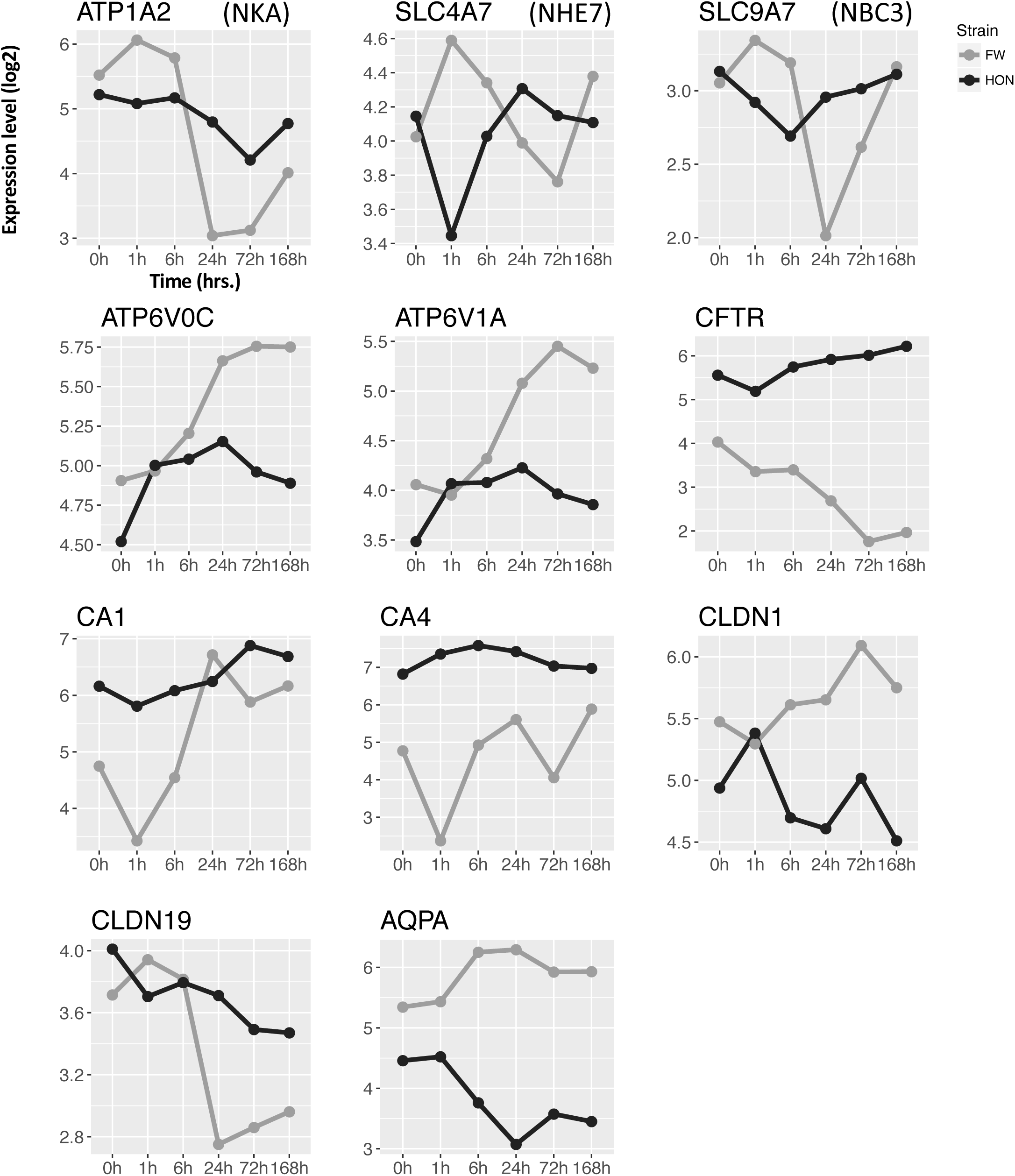
Log2 expression levels for genes typically expressed in ionocytes, including sodium/potassium-transporting ATPase subunit alpha-2 (ATP1A2, or NKA), sodium bicarbonate cotransporter 3 (SLC4A7, or NBC3), sodium/hydrogen exchanger 7 (SLC9A7, or NHE7), V-type proton ATPase 16 kDa proteolipid subunit (ATP6V0C), V-type proton ATPase catalytic subunit A (ATP6V1A), cystic fibrosis transmembrane conductance regulator (CFTR), carbonic anhydrase 1 (CA1), carbonic anhydrase 4 (CA4), claudin-1 (CLDN1), claudin-19 (CLDN19), and aquaporin FA-CHIP (AQPA), after time in air (hours) and in pre-emersion controls (0h) for HON (black) and FW (grey) fish.

**Figure S15.**
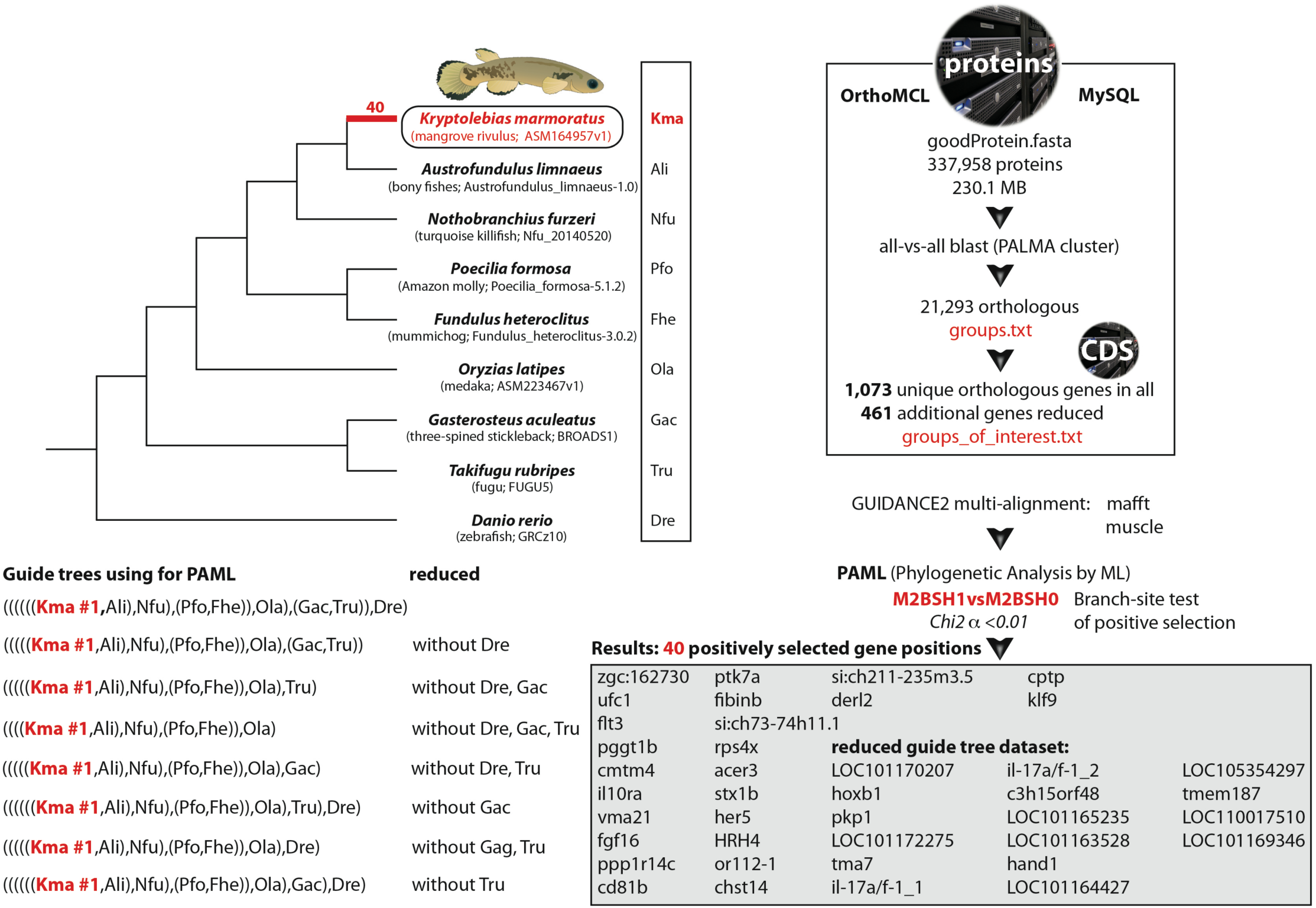
Workflow for detecting genes evolving by positive selection in amphibious *K. marmoratus* compared to other non-amphibious fish.

